# Glutamate indicators with increased sensitivity and tailored deactivation rates

**DOI:** 10.1101/2025.03.20.643984

**Authors:** Abhi Aggarwal, Adrian Negrean, Yang Chen, Rishyashring Iyer, Daniel Reep, Anyi Liu, Anirudh Palutla, Michael E. Xie, Bryan J. MacLennan, Kenta M. Hagihara, Lucas W. Kinsey, Julianna L. Sun, Pantong Yao, Jihong Zheng, Arthur Tsang, Getahun Tsegaye, Yonghai Zhang, Ronak H. Patel, Benjamin J. Arthur, Julien Hiblot, Philipp Leippe, Miroslaw Tarnawski, Jonathan S. Marvin, Jason D. Vevea, Srinivas C. Turaga, Alison G. Tebo, Matteo Carandini, L. Federico Rossi, David Kleinfeld, Arthur Konnerth, Karel Svoboda, Glenn C. Turner, Jeremy Hasseman, Kaspar Podgorski

## Abstract

Identifying the input-output operations of neurons requires measurements of synaptic transmission simultaneously at many of a neuron’s thousands of inputs in the intact brain. To facilitate this goal, we engineered and screened 3365 variants of the fluorescent protein glutamate indicator iGluSnFR3 in neuron culture, and selected variants in the mouse visual cortex. Two variants have high sensitivity, fast activation (< 2 ms) and deactivation times tailored for recording large populations of synapses (iGluSnFR4s, 153 ms) or rapid dynamics (iGluSnFR4f, 26 ms). By imaging action-potential evoked signals on axons and visually-evoked signals on dendritic spines, we show that iGluSnFR4s/4f primarily detect local synaptic glutamate with single-vesicle sensitivity. The indicators detect a wide range of naturalistic synaptic transmission, including in the vibrissal cortex layer 4 and in hippocampal CA1 dendrites. iGluSnFR4 increases the sensitivity and scale (4s) or speed (4f) of tracking information flow in neural networks *in vivo*.

## Introduction

Neurons process information by combining and transforming signals arriving at their many synaptic inputs^1–3^. A major goal of brain research is therefore to monitor the activity of input synapses and neuronal output in the intact brain. Fluorescent calcium indicators^4^, voltage indicators^5–7^, and extracellular electrophysiology^8^ are routinely used to record outputs in large populations of neurons^9^. In contrast, technologies to record from large populations of neurons’ synaptic inputs do not yet exist.

The large majority of vertebrate central synapses release the neurotransmitter glutamate^3^. One action potential (AP) typically releases zero or one vesicles, freeing a few thousand glutamate molecules, which are cleared from the synaptic cleft in less than one millisecond^10^. For comparison, one AP triggers influx of over 10^5^ calcium ions into a typical neuronal cell body, which are cleared in 20 milliseconds^11^. The small number of glutamate molecules and their short residence time make optical measurements of synaptic release extremely challenging, especially when recording from many synapses at once *in vivo*. Addressing this challenge is necessary to reveal fundamental principles of neuronal computation, such as whether the nonlinearities that implement computations primarily occur in the soma or dendrites^1,12^, and what patterns of synaptic input drive neuronal firing and plasticity *in vivo*^2^. Neurotransmitter recordings can also be used to study synaptic plasticity^13–17^, neural connectivity^18^, neurological disorders^19–21^, and the molecular mechanisms and pharmacology of synaptic transmission^17,22–24^.

Fluorescent protein neurotransmitter indicators are fusions between a binding domain and a fluorescent protein domain, displayed on the cell surface^25^. Conformational changes upon ligand binding modulate the indicator fluorescence, which can be read out at high resolution with a microscope. The fluorescent protein glutamate indicator iGluSnFR can detect release of individual synaptic vesicles under favorable conditions^13,15,22^, but even for the highest signal-to-noise-ratio (SNR) variant iGluSnFR3^26^, SNR *in vivo* is often insufficient to record from more than a few dozen synapses at once. SNR decreases when imaging more synapses because limited total excitation power must be divided across synapses^27,28^. Brighter, more sensitive indicators are required for measurements from large groups of individual synapses.

Additional constraints arise from the finite voxel rates of microscopes, imposing a tradeoff between sampling rate and number of synapses recorded. For example, using conventional two-photon microscopy, up to a few dozen of a neuron’s synapses can be addressed at the ∼ 100 Hz frame rate required to resolve iGluSnFR3 glutamate transients, which decay with a time constant below 30 ms^26^. Variants with slower deactivation kinetics would enable imaging with lower frame rates and thus access larger populations of synapses. In contrast, variants with fast deactivation kinetics would allow more precise monitoring of rapid synaptic dynamics^29,30^, albeit limited to smaller imaging volumes. Regardless of deactivation rate, fast activation kinetics are needed for precise measurements of glutamate release times^27,31^.

Here we engineered two highly sensitive, fast-activating iGluSnFR variants with fast or slow deactivation. A large library of rationally-targeted mutations was screened *in vitro* for improved brightness, kinetics and sensitivity. Seven variants were further tested using two-photon imaging in the mouse visual cortex. We selected two variants, iGluSnFR4f (fast deactivation) and iGluSnFR4s (slow deactivation), with significantly improved sensitivity and brightness. We highlight their advantages in experiments involving synaptic imaging in the visual cortex, somatosensory cortex, and hippocampus, and fiber photometry in the midbrain.

## Results

### Cultured neuron field stimulation screen

Previous research identified many sites that influence iGluSnFR function^26,29,30,32^, but the large sequence space across these sites has not been well explored. We conducted saturating mutagenesis at 41 previously-identified sites in two iGluSnFR3 variants (iGluSnFR3.v857 and iGluSnFR3.v867), generating a total of 1,640 variants (Figs 1a-b). Primary cortical cultures from rats (DIV14) were transduced with these, and their responses to field stimulation were imaged^33^. (Figs 1c-d). Baseline fluorescence brightness (F_0_), peak fractional response (ΔF/F_0_), and rise and decay times (T_on_ and T_off_) were measured (Figs. 1e,f,g, S1).

**Figure 1:**
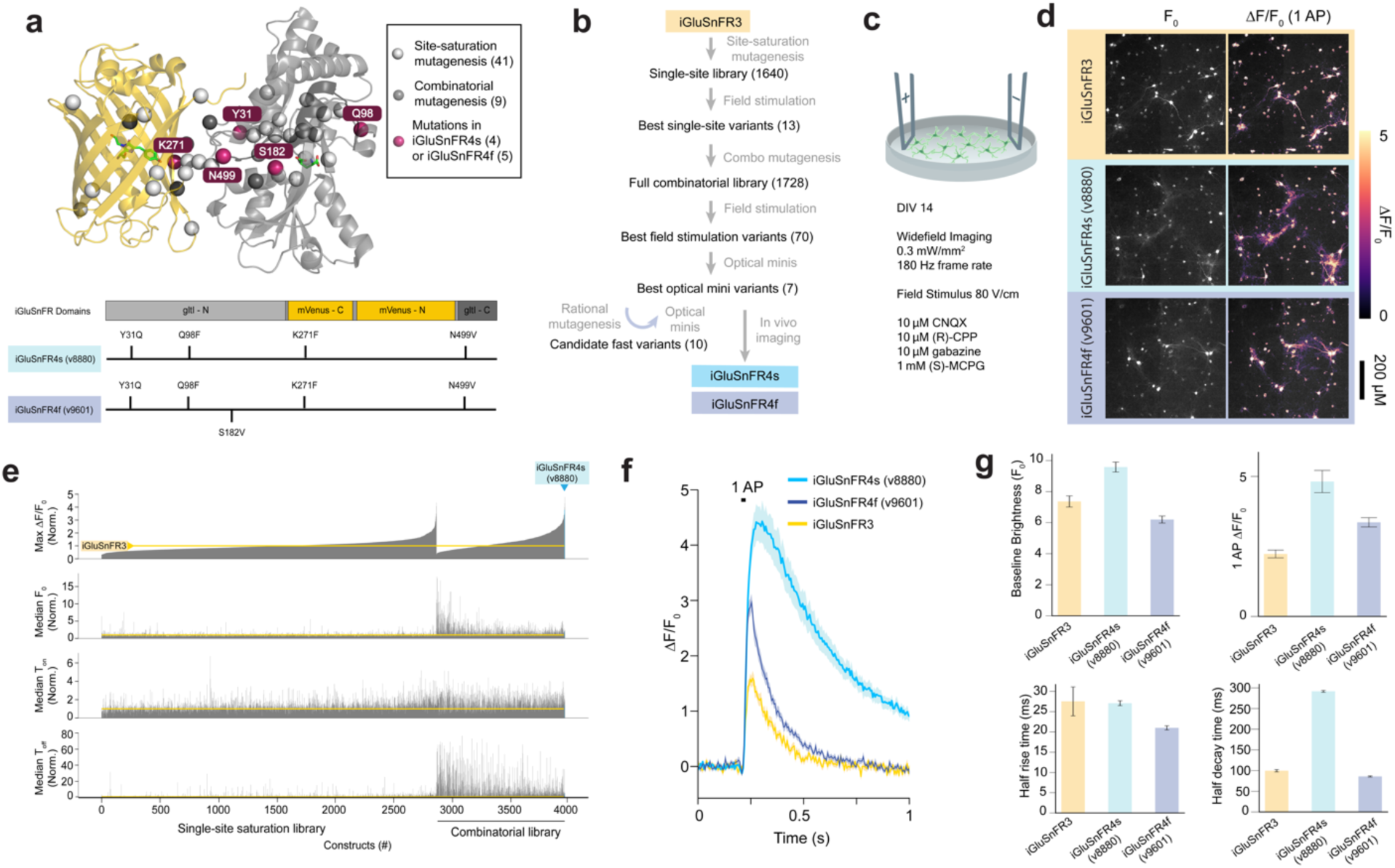
Screening iGluSnFR variants in primary neuronal culture. a) Top, location of sites chosen for mutagenesis mapped onto the iGluSnFR3 structure. Light gray spheres, sites selected for saturation mutagenesis; dark gray sphere, sites included in full combinatorial mutagenesis; red spheres, sites included in iGluSnFR4s or 4f. Bottom, mutations in iGluSnFR4s and 4f. **b)** Screening steps. **c)** Primary neuron cultures expressing iGluSnFR variants received a brief electric field stimulus to evoke a single action potential. **d)** Images of cultures expressing iGluSnFR3, iGluSnFR4s, and iGluSnFR4f at baseline ( F_0,_ left), and peak fluorescence change (ΔF/F_0_, right) following a single field stimulus. **e)** Measured ΔF/F_0,_ F_0_, rise time (T_on_) and decay time (T_off_), normalized to within-plate iGluSnFR3 controls, ordered by ΔF/F_0_ for significantly-responding constructs in the single-site and combinatorial libraries. N = 4-16 wells/variant. **f)** Mean ΔF/F_0_ fluorescence traces in the field stimulation assay for iGluSnFR3, iGluSnFR4f and 4s. g) Summary of F_0_, Peak ΔF/F_0_, T_on_, and T_off_ of the three variants. N (wells) = 66, 74, 73. Errorbars denote SEM.

From this screen, we selected 12 point mutations across 9 sites (Y31Q, Y31E, Q34A, Q98F, A185N, T254R, K271F, K271G, H273E, Q418S, N499L, and N499V) that exhibited superior performance in at least one parameter (F_0_, ΔF/F_0_, T_on_, T_off_). We then constructed a complete combinatorial library comprising 1,728 variants on the iGluSnFR3.v857 background (hereafter referred to as iGluSnFR3). Of these, 1,392 variants were successfully expressed and exhibited detectable responses to field stimuli, and were included in further analyses.

We quantified effects of each mutation and their interactions using a generalized linear model (GLM) (Supplementary Fig. 1a). Incorporating pairwise interactions into the model enhanced the cross-validated variance explained for all response variables, indicating the presence of epistatic interactions (Supplementary Fig. 1b–e). We resolved a high-resolution crystal structure of iGluSnFR3 (PDB: 9FBU), allowing us to map mutation effects and interactions to physical positions. The magnitude of pairwise interactions was inversely correlated with distance (r=-0.28, p=5.34e-06; Supplementary Fig. 1f). Although local in physical space, many of the strongest interactions spanned domains of the fusion protein (Supplementary Fig. 1g).

### Optical mini screen

We selected 70 variants from the combinatorial screen and tested them by imaging spontaneous synaptic glutamate release (‘optical minis’) in cultured neurons silenced with tetrodotoxin (TTX). Optical minis are caused by asynchronous release of individual synaptic vesicles^22,26,34,35^, producing events with a much smaller spatial span and therefore faster diffusion-limited kinetics than field-evoked responses (Fig 2a-c). We reasoned that optical mini imaging would be well-suited to identify variants with fast decay kinetics and improved SNR for synaptic glutamate dynamics. Many variants showed higher SNR than iGluSnFR3, with the top-performing variant, v8880 (iGluSnFR3+Y31Q,Q98F,K271F,N499V), having 4.7-fold higher SNR and 1.2-fold slower decay rate (Fig. 2d; Supplementary Videos 1). Variants in the combinatorial screen and mini screens had slower-decaying signals than iGluSnFR3 (Fig 1e, 2d), suggesting that the field stimulation screen favored slower indicators. To create high-SNR variants with fast decay kinetics, we introduced five additional mutations known for short decay rates (S70A, S70T, S182L, S182V, or Y209F)^32,30,29,36^ onto two high-sensitivity variants, v8880 and v8376. These ten variants (v9598 through v9607) were further characterized with optical mini recordings. The best-performing of these, v9601 (v8880+A182V), showed 2.1-fold higher SNR and 3.2-fold faster decay versus iGluSnFR3 (Fig 2d; Supplementary Video 1).

**Figure 2:**
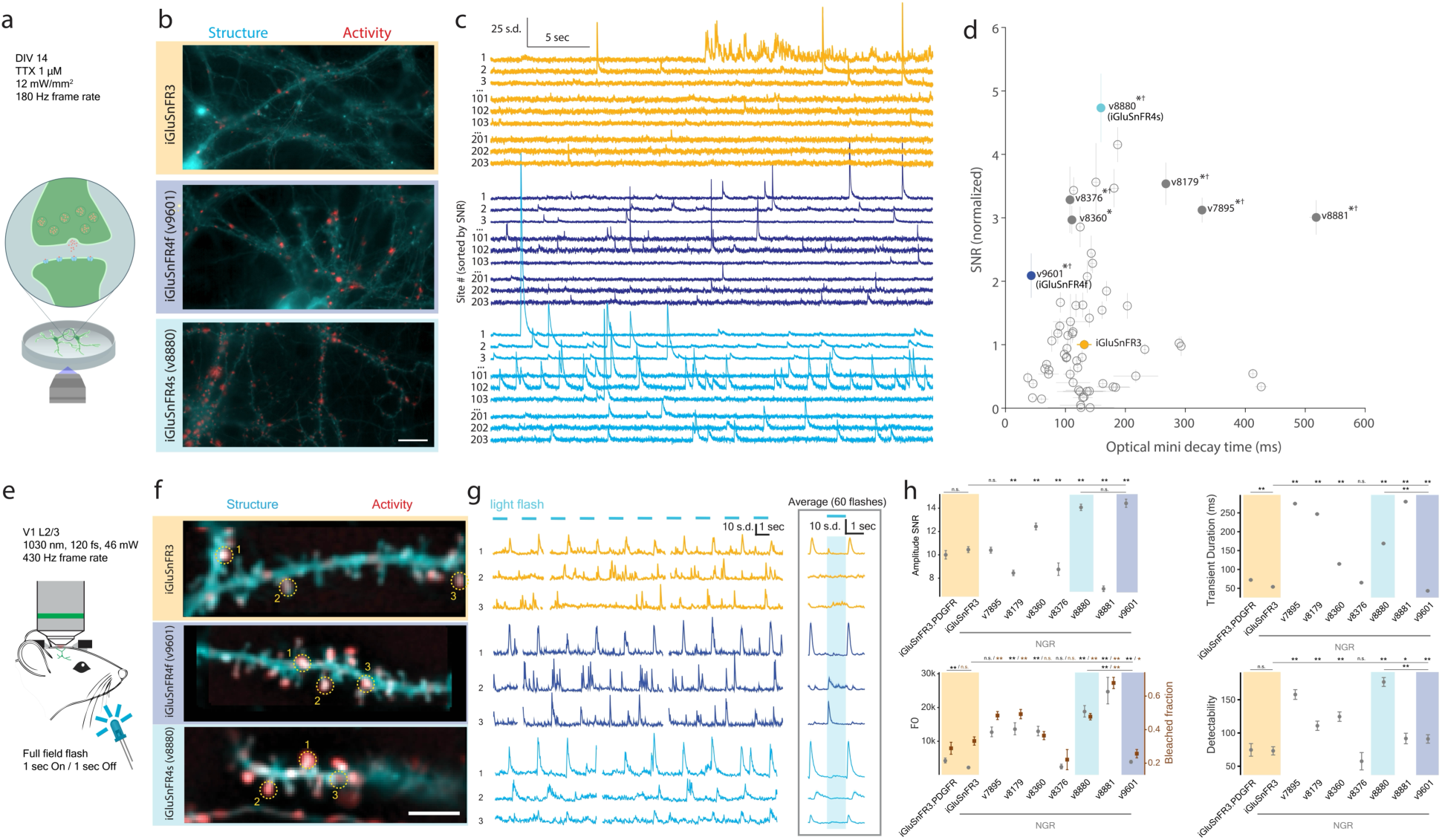
Synapse-resolved screening in cultured neurons and the visual cortex *in vivo*. a) iGluSnFR variants underwent high-speed, high-resolution widefield imaging of spontaneous release events (optical minis) in primary neuronal cultures silenced with tetrodotoxin (TTX). **b)** Representative structural (cyan; gamma 0.5) and pixel-wise activity (normalized skewness, red) images for iGluSnFR3, v9601 (iGluSnFR4f), and v8880 (iGluSnFR4s), on matching color scales across panels for both channels. Scale bar: 20 µm. **c)** ΔF traces for individual sites for each variant, each normalized to its standard deviation (s.d.), and ordered by SNR. Sites ranked 1–3, 101-103, and 201-203 are shown for each. **d)** SNR and decay time (Mean+/-SEM across wells) of all variants in optical mini screen. Filled circles denote variants selected for in vivo characterization. n=12-16 wells (iGluSnFR4 variants); n = 99 wells (iGluSnFR3). †: p<0.05 decay time vs iGluSnFR3; *: p<0.05 SNR vs iGluSnFR3. Two-tailed Mann-Whitney test. **e)** Dendrites of sparsely-labeled layer 2/3 neurons in V1 were recorded with 2P imaging while presenting a full field light flash stimulus. **f)** Representative structural images (cyan; gamma 0.5) and pixel-wise activity (red) for iGluSnFR3, v9601 (iGluSnFR4f), and v8880 (iGluSnFR4s), on matching color scales across panels for both channels. Scale bar: 10 µm. **g)** Representative traces (left), and stimulus triggered averages (right) for ROIs shown in (f). Gaps in traces correspond to frames discarded due to movement. **h)** Summaries of amplitude SNR, transient duration, F0, bleached fraction (larger values denote more bleaching), and detectability for the screened variants. Except for transient duration which was averaged directly, data points and errorbars are GLM fits and S.E. CIs accounting for covariates of expression time and imaging depth. N (distinct branches) = 15 iGluSnFR3.PDGFR, 52 iGluSnFR3, 19 v7895, 17 v8179, 31 v8360, 40 v8880, 12 v8881, 25 v9601).

### Mouse visual cortex screen and selection of iGluSnFR4s and 4f

We selected 7 variants for *in vivo* testing (Fig 2e-h). We used the NGR signal sequence^26,37^ for membrane display, and compared against iGluSnFR3 in both NGR and PDGFR vectors as controls. We sparsely labeled neurons in primary visual cortex (V1) by coinfection of adeno-associated virus (AAV) expressing Cre-dependent iGluSnFR and low titers of AAV expressing Cre. First, we imaged layer 2/3 pyramidal neuron dendrites while presenting periodic full-field light-flash visual stimuli (1s on, 1s off). Fluorescence transients were extracted using a non-negative matrix factorization based algorithm. Amplitude SNR, detectability (an SNR measure that accounts for transient duration), F_0_, and photobleaching were quantified using a GLM to control for imaging depth and expression time (Fig 2h; Methods). Photobleaching was highly correlated with F_0_ (r^2^=0.86, p=2.9e-4), suggesting that variations in bleaching across variants were due largely to differences in per-molecule excitation rate at baseline. v8880 and v9601 showed the highest SNR among variants tested, consistent with the optical mini screen. Relative transient durations *in vivo* were also consistent with the optical mini screen, with v9601 the fastest variant tested. Despite shorter transients, v9601 exhibited higher detectability than iGluSnFR3.

Dendritic spines are specialized protrusions that receive most glutamatergic inputs onto pyramidal neurons, with addition, retraction, and morphological changes regulated by activity^38^. We tested whether expression of iGluSnFR variants affects spine survival. We did not see differences in iGluSnFR-expressing neurons versus neurons expressing membrane-tagged EGFP (Supplementary Fig. 2).

Based on these assays, we selected v8880 and v9601 as the best-performing slow- and fast- decay variants, respectively, naming them iGluSnFR4s and iGluSnFR4f. We characterized both variants in purified soluble protein (Supplementary Table 1, Supplementary Figs. 3-6) and on cultured neurons (Supplementary Fig. 7,8). Purified protein measurements revealed strong modulation of both extinction and fluorescence quantum yield upon glutamate binding and high selectivity over other neurotransmitters and related amino acids. In neuronal culture, iGluSnFR4s and iGluSnFR4f exhibit higher single-AP ΔF/F₀ than iGluSnFR3 (Fig. 1d–g), and higher (4s) or lower (4f) on-membrane affinity (Supplementary Fig. 7). iGluSnFR4s has similar rise and much slower decay versus iGluSnFR3, while 4f has faster rise and decay (Fig. 1i). As a result, iGluSnFR4f better follows rapid synaptic release in culture than iGluSnFR3 (Supplementary Fig. 8).

### iGluSnFR4 detects synaptic glutamate in the visual cortex

To assess *in vivo* performance, we first expressed iGluSnFR3, iGluSnFR4s and iGluSnFR4f sparsely in neurons in primary visual cortex (V1) using single-cell electroporation. We then used loose cell-attached recording and two-photon imaging to characterize single-AP evoked glutamate transients on axonal boutons of the recorded neurons (Fig. 3a,b). All three indicators exhibited fluorescence transients time-locked with somatically-recorded APs (Fig. 3b,c). We computed spike-triggered averages (STAs) (Fig. 3c), which were well described by a fast rise and slower exponential decay. iGluSnFR4f and 4s showed significantly larger 1AP-evoked amplitudes than iGluSnFR3 (Fig 3d). All three indicators showed rise times less than 2 ms (Fig 3d). iGluSnFR4f showed faster (median 25.9 ms, IQR 24.2-27.6 ms) and iGluSnFR4s much slower (median 152.7 ms, IQR 134.9-163.5 ms) decay time constants than iGluSnFR3 (median 29.1 ms, IQR 25.6-32.6 ms) (Fig. 3d). The iGluSnFR4 variants each showed increased detectability of single APs over iGluSnFR3 (Supplementary Fig. 9).

**Figure 3:**
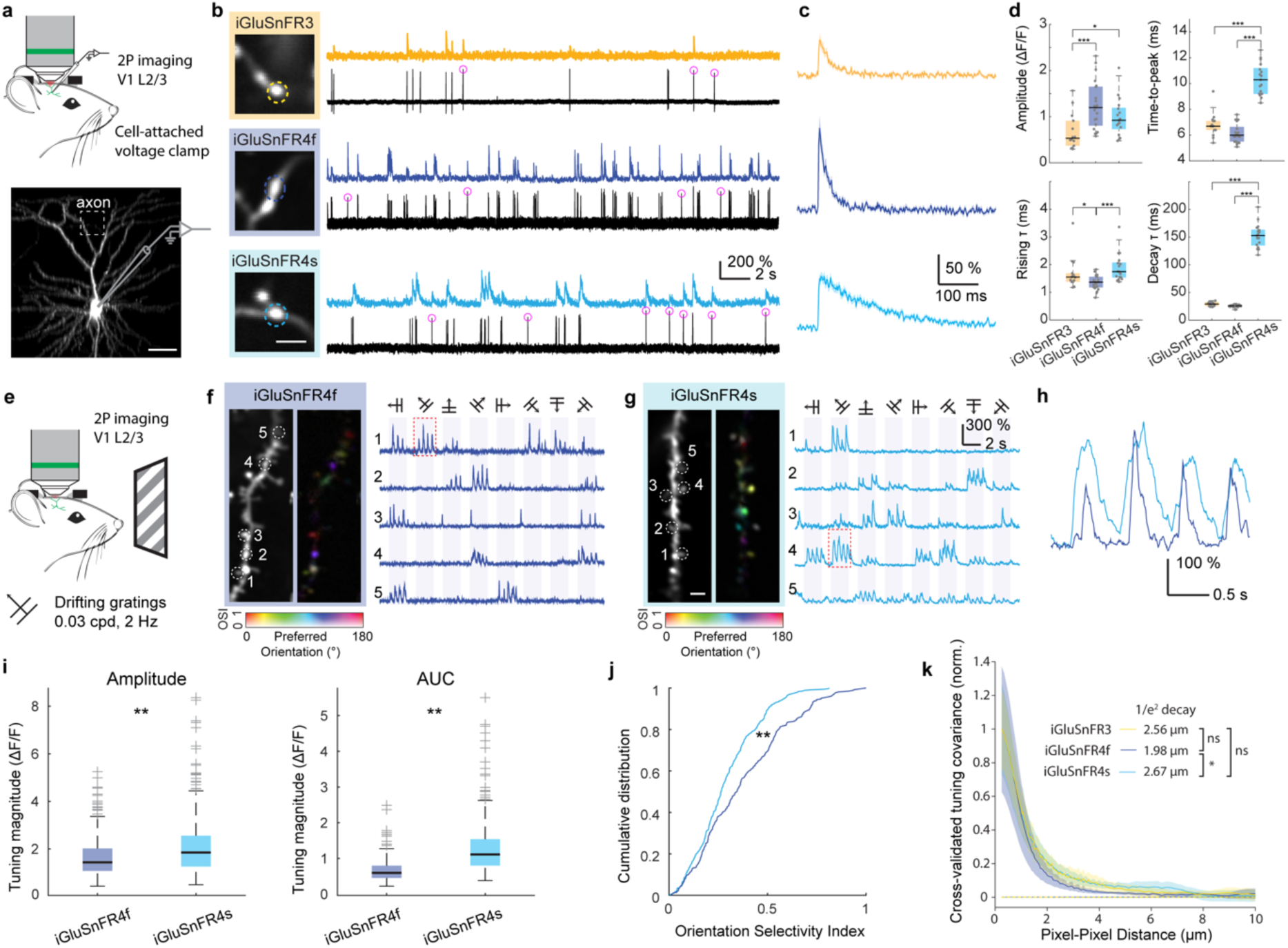
Characterization of iGluSnFR4 in the mouse visual cortex. a) Layer 2/3 neurons in V1 were transduced with plasmids expressing iGluSnFR variants using single-cell electroporation. Following expression, simultaneous axonal imaging and loose-seal, cell-attached recordings were performed. Scale bar, 50 μm. **b)** Axonal glutamate signals and somatic spiking for neurons expressing iGluSnFR3, iGluSnFR4f, and iGluSnFR4s. Pink circles indicate single APs. Scale bar, 2 μm. **c)** Spike-triggered averages for 1AP-evoked axonal glutamate signals. Shading denotes SEM. N= 46 iGluSnFR3, 24 iGluSnFR4f, 58 iGluSnFR4s isolated spikes. **d)** Amplitude, time-to-peak, full-width half-maximum (FWHM), rising τ, and decay τ for the 1AP glutamate transients from iGluSnFR3 (n = 14 boutons from 4 cells), iGluSnFR4f (n = 21 boutons from 4 cells) and iGluSnFR4s (n = 17 boutons from 4 cells). *p < 0.05; ***p < 0.001; Kruskal-Wallis test with Dunn’s test for multiple comparisons. **e)** L2/3 dendrites were imaged while stimulating the contralateral eye with oriented drifting gratings. **f)** Two-photon image of a dendritic segment from an iGluSnFR4f labeled L2/3 neuron and a pixel-wise tuning map (left). Representative single-trial ΔF/F_0_ traces from denoted ROI are shown on the right. **g)** Same as above but for iGluSnFR4s. Scale bar, 2 μm. **h)** Expanded traces from the red boxes. **i)** Tuning magnitude for iGluSnFR4f and iGluSnFR4s computed from the response amplitude (left, n = 324 ROIs for iGluSnFR4f and n = 349 ROIs for iGluSnFR4s from 4 cells each, P = 9.6x10-9, Wilcoxon rank sum test) or the area under the curve (right, n = 267 ROIs for iGluSnFR4f and n = 348 ROIs for iGluSnFR4s from 4 cells each. P = 8.4x10-49, Wilcoxon rank sum test). Boxplots denote median and interquartile range, outliers beyond 1.5 interquartile plotted separately. **j)** Distribution of orientation selectivity index for iGluSnFR4s and iGluSnFR4f (n = 267 ROIs for iGluSnFR4f and n = 348 ROIs for iGluSnFR4s from 4 cells each, P = 4.8x10-6, Kolmogorov–Smirnov test). **k)** Cross-validated covariances for pairs of labeled pixels at various distances, indicative of the spatial spread of iGluSnFR signals around release sites, for iGluSnFR3, 4s, and 4f. Dashed curves denote covariances calculated over unlabeled pixels. N = 8 FOVs per variant. Errorbars denote bootstrapped 95% ci. Mean distances to 1/e^2^ of peak labeled pixel covariance are shown. *: p<0.05, bootstrap test.

Next, we compared the ability of the iGluSnFR variants to resolve signals at individual synapses. Cortical neuropil contains approximately one glutamatergic synapse per cubic micrometer^39,40^. The kinetic properties of indicators shape the spatial extent of fluorescence signals^26^. Indicators that saturate near the site of release, where glutamate concentrations are high, result in signals with larger spatial extent that produce measurement crosstalk between nearby synapses.

To assess spatial extent and crosstalk, we imaged dendritic responses to directional drifting grating stimuli in V1 (Fig 3e). Synaptic inputs to V1 neurons have diverse preferences for grating orientation, direction, and phase, exhibiting little spatial organization over micrometer scales^41^. Spatial crosstalk would blend signals with different tuning, reducing measured selectivity. Both iGluSnFR4f and 4s reported high signal-to-noise, orientation-tuned responses localized to dendritic spines (Fig 3f-j). We observed clear responses for each cycle of the grating stimuli from many distinct sites. iGluSnFR4f responses were sharper in time (Fig. 3f-h), whereas iGluSnFR4s responses have larger peak amplitudes and integrated areas (Fig. 3i). The percentage of tuned responses reported by iGluSnFR4s was lower than 4f (Fig. 3j), suggesting slightly more crosstalk with the slower, higher-affinity indicator. We next measured the spatial extent of signals using cross-trial tuning covariances. Because nearby synapses have largely uncorrelated tuning, measured covariance in tuning between pairs of pixels (calculated from non-overlapping trials; Methods) reflects the spatial spread of signals arising from a synapse^42^. iGluSnFR4f showed a slightly narrower covariance than iGluSnFR4s, although neither differed significantly from iGluSnFR3 (Fig. 3k). These results show that iGluSnFR4f and 4s report signals from individual synapses with high specificity similar to iGluSnFR3, although 4f has slightly higher specificity than 4s.

### Frequency response characterization in barrel cortex layer 4

Rodents sense the world by rhythmically sweeping their vibrissae (whiskers) over object features at approximately 15 Hz^43^. Signals from the vibrissa follicles ascend to the primary somatosensory cortex, vS1, via the ventral posteromedial (VPM) thalamus. To characterize stimulus-evoked glutamatergic signals across frequencies, we expressed iGluSnFR3, 4s and 4f by injecting AAVs into VPM, then recorded from thalamocortical axons 350-400 μm deep in vS1. Imaging was done with an adaptive-optics two-photon (AO-2P) microscope in head-fixed mice during rhythmic air-puff stimulation (Fig 4a-d) and voluntary whisking (Fig 4e-i).

**Figure 4:**
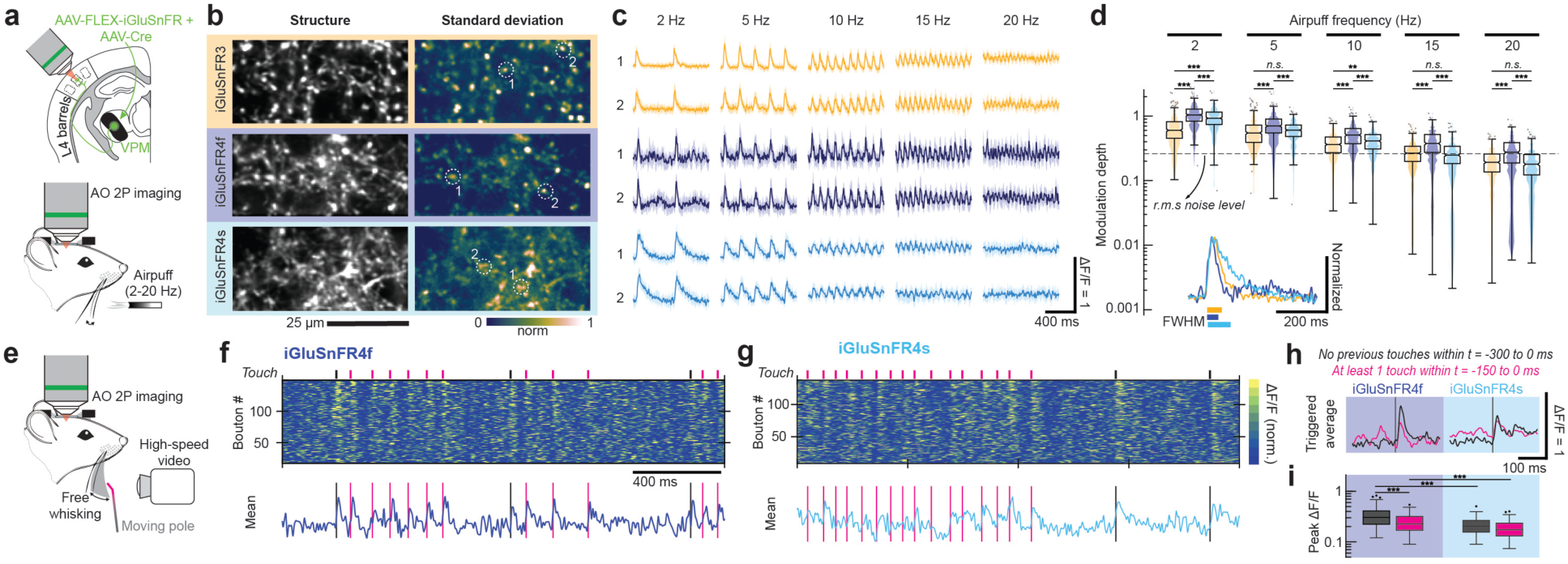
Imaging rapid dynamics in thalamocortical axons using AQ-2P scanning microscopy. a) iGluSnFR variants were expressed by AAV injection in VPM and thalamocortical boutons in barrel cortex recorded with AO-2P imaging. Awake mice received rhythmic whisker airpuff stimulation during high-speed AO-2P imaging. **b)** (left) Average image of thalamocortical axons in L4 labeled by the iGluSnFR variants. (right) Normalized pixelwise standard deviation across 1-s averaged-epochs from 60 trials at 5 Hz airpuff stimulus. **c)** Mean responses to the indicated airpuff frequencies for boutons marked in (**c**); shaded area indicates the 25th and 75th percentiles (n = 20 trials). **d)** Comparison of iGluSnFR variant response amplitudes in thalamocortical boutons at different stimulation frequencies during a 4-second airpuff stimulus: iGluSnFR3 (three mice, 431 boutons), iGluSnFR4f (two mice, 407 boutons), iGluSnFR4s (three mice, 409 boutons). The dashed line indicates root-mean-square (r.m.s.) noise level for iGluSnFR3. ***p<0.0001, **p<0.001; two-sample t-test with unequal variances. Inset, normalized airpuff-triggered average for a single bouton for each of the iGluSnFR variants, mean over 100 stimulus pulses. Extent above half-maximum is plotted below. **e)** Awake mice were allowed to whisk freely against a moving pole during high-speed AO2P imaging of thalamocortical boutons and high-speed whisker videography to record touch events **f)** (top) Chart showing ΔF/F_0_ traces of individual boutons and (bottom) average ΔF/F_0_ traces of the 10 most responsive boutons over a 2-second period from a single field labelled with iGluSnFR4f. The black and pink lines mark the instances of the two classes of pole touch events in (i). **g)** As in (f), for iGluSnFR4s. **h)** Touch-triggered mean responses. Black curves indicate responses from touch events with no preceding touches within 300 ms (21 events iGluSnFR4f, 27 events iGluSnFR4s). Pink curves indicate responses with at least one touch event in the preceding 150 ms (67 events iGluSnFR4f, 88 events iGluSnFR4s). **i)** Distribution of peak responses from the two categories, for all boutons. (iGluSnFR4f: two mice, 127 boutons, iGluSnFR4s: two mice, 148 boutons), ***p<0.0005.

We observed rapid-onset responses, time-locked to rhythmic stimulation from 2-30 Hz with all indicators (809 boutons, 7 mice, Fig. 4c, Supplementary Video 2). The modulation depth of iGluSnFR signals decreased with stimulus frequency, in a manner that was consistent with each variant’s finite response bandwidth (Fig. 4c). Nevertheless, iGluSnFR4f was highest-modulated at all frequencies (Fig. 4d). iGluSnFR4f responses were detected up to 20 Hz, above the natural whisking frequency of mice.

We next characterized iGluSnFR4 touch responses during free-whisking against a pole while recording touch events with a high-speed camera (Fig. 4e). The pole was moved randomly to add variability to inter-touch intervals. We compared two categories of touch events: ones with no touches in the preceding 300 ms and ones with at least 1 touch in the preceding 150 ms. While iGluSnFR4s consistently captured the former category, responses to rapidly recurring touch events were poorly resolved due to its slow decay time, and iGluSnFR4f produced resolvable responses for both categories (Fig. 4f,g). On average, iGluSnFR4f produced higher-amplitude responses for both categories (Fig. 4h,i) and is well-suited for monitoring inputs to somatosensory cortex in rodents.

### Video rate imaging of iGluSnFR4s in layer 5 neuron tuft dendrites

Full-frame (e.g. 512 scanlines) resonant scanning two-photon imaging is limited to ∼30 Hz frame rate, too slow to adequately sample iGluSnFR3 transients. We evaluated the sensitivity of iGluSnFR4s for large-scale video-rate recording by imaging tuft dendrites of AAV-transfected Layer 5 neurons in mouse V1 during presentation of brief sparse noise stimuli (Supplementary Fig. 10a), which drive weaker responses than drifting gratings. Each dendrite showed sites responsive to stimuli localized within the visual field, estimated by fitting a cross-validated spatiotemporal receptive field (stRF) (Supplementary Fig. 10b-g). We considered sites responsive whose stRF explained more variance than expected by chance and over 10% of total variance. iGluSnFR4 yielded larger responses, higher SNR, more variance explained per ROI, and many more responsive sites per dendrite than iGluSnFR3 under video-rate imaging (Supplementary Fig. 10j-m);

### High-sensitivity photometry with iGluSnFR4s

Fiber photometry, which involves delivering excitation light and collecting resulting fluorescence via an optical fiber without forming an image, has been broadly adopted for neuroscience measurements in deep regions^44,45^. Because photometry integrates signals from all labeled membranes it benefits from high-affinity indicators, which capture more ligand farther from release sites. Photometry also benefits from slower indicators because these generate more time-integrated photons per release event. The large volumes sampled in photometry recordings tend to result in less-synchronized, slower signals than two-photon imaging, reducing the benefits of very fast indicators. For these reasons, iGluSnFR3 shows similar-amplitude photometry responses to the slower, higher-affinity variant SF-iGluSnFR.A184S^45^, despite producing much larger signals in high-resolution imaging^26^. We reasoned that iGluSnFR4s would be well-suited for photometry. We expressed iGluSnFR4s and either iGluSnFR3 or SF-iGluSnFR in ventral tegmental area (VTA) GABAergic neurons, paired across brain hemispheres (Fig. 5a). We recorded from the paired indicators while presenting water-restricted mice with water rewards (Fig. 5b). iGluSnFR4s produced responses with larger amplitude than iGluSnFR3 (5.62 ± 0.42 vs 1.56 ± 0.16) or SF-iGluSnFR (3.72 ± 0.92 vs 1.54 ± 0.44) in the paired measurements (Fig. 5c-d).

**Figure 5:**
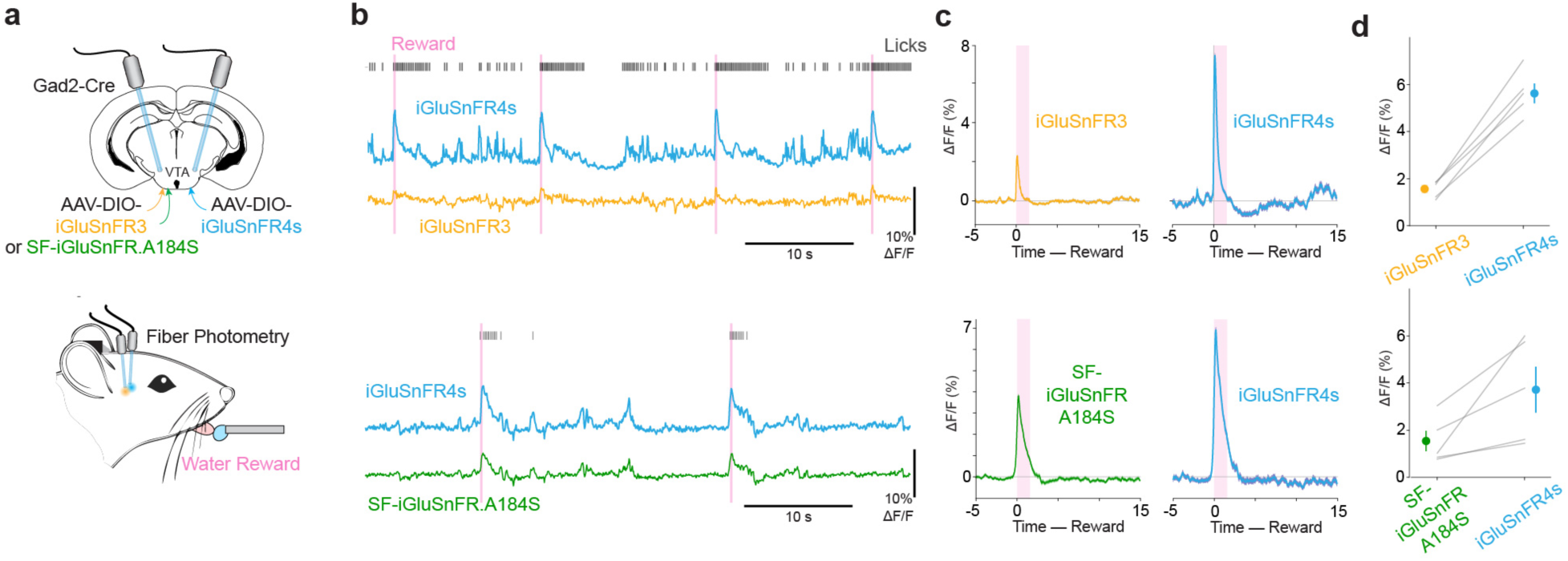
Deep-brain fiber photometry with iGluSnFR4s. a) iGluSnFR4s and either iGluSnFR3 or SF-iGluSnFR.A184S were expressed virally, and fibers implanted, in the ventral tegmental areas of opposite hemispheres. Photometry from both hemispheres was performed in mice receiving periodic water rewards. **b)** Example recordings demonstrating responses at stimulus time and during licks. **c)** Trial averaged ΔF/F_0_ traces of the three indicatos aligned to reward delivery. Shaded region shows the time window used to quantify mean ΔF/F_0_ in (d). **d)** Mean and SEM post-reward ΔF/F_0_ for the paired-hemisphere recordings. P = 4.0 × 10^-4^ (3 vs 4s); P = 0.0516 (SF vs 4s); paired t-test

## Discussion

Here we developed new iGluSnFR variants with improved sensitivity and faster (iGluSnFR4f: 27 ms) or slower (iGluSnFR4s: 126 ms) deactivation times. As with calcium indicators^5,46^, fast- and slow- deactivating iGluSnFRs each have distinct and valuable applications. iGluSnFR4f better follows rapid inter-release-intervals and shows higher spatial specificity (Figs. 3k,4,S8). iGluSnFR4s produces larger time-integrated photon counts and SNR in two-photon recordings, greatly increased signals under slower (video-rate) imaging and greatly increased signals in fiber photometry (Figs. 2,3,5, S9). Despite its slower decay, iGluSnFR4s shows fast rise kinetics (<2 ms), a combination that can enhance detectability while allowing precise timing inference from transients’ rising phase^31^ (Figs. 2h,3d).

Indicator SNR is critical for observing single-vesicle release^13,15,22,26^, recording from larger synaptic populations^47^, and other challenging applications. Recording from larger populations is needed to study the potentially complex but poorly understood input-output operations of individual neurons, which integrate input from thousands of excitatory synapses on milliseconds timescales^1,2^.

iGluSnFR4 was evolved using assays in neuron culture and *in vivo* designed to select for responses to individual release events, such as 1 AP field stimulation and optical mini imaging. The relative performance of tested variants was consistent across these assays. However, both iGluSnFR4 variants show reduced dynamic range to saturating ligand in soluble protein, a commonly-used screening assay. This highlights the importance of screening with assays that match intended applications. Our combinatorial screen densely sampled application-relevant functional properties over the high-dimensional sequence space of a set of 12 mutations that each have strong individual effects. We identified numerous nonlinear interactions, which may be useful for indicator engineering beyond this work, for example to train and test models that predict complex protein sequence-to-function relationships.

We demonstrated wide applicability of iGluSnFR4f/4s including *in vivo* recordings of axons and dendrites, and cell types from layers 2-5 of neocortex, thalamus, hippocampus (Supplementary Fig. 11), and GABAergic midbrain neurons.

We performed tests under both the PDGFR (Figs 2f-h, 3a-j, 4,5) and NGR (Figs 2f-h, 3k,S7,S8) membrane display vectors. The NGR vector matched or outperformed PDGFR under side-by-side tests *in vivo*, and greatly outperformed PDGFR in culture^26,37^. We recommend the NGR vector for most applications, particularly in culture.

In conclusion, iGluSnFR4s and 4f address critical challenges for synaptic imaging, particularly *in vivo* recordings from populations of individual synaptic inputs and rapid synaptic dynamics. These tools will enable new studies in neuronal computation, synaptic physiology, and mechanisms underlying brain disorders.

## Supplementary Figures

**Supplementary Figure 1:**
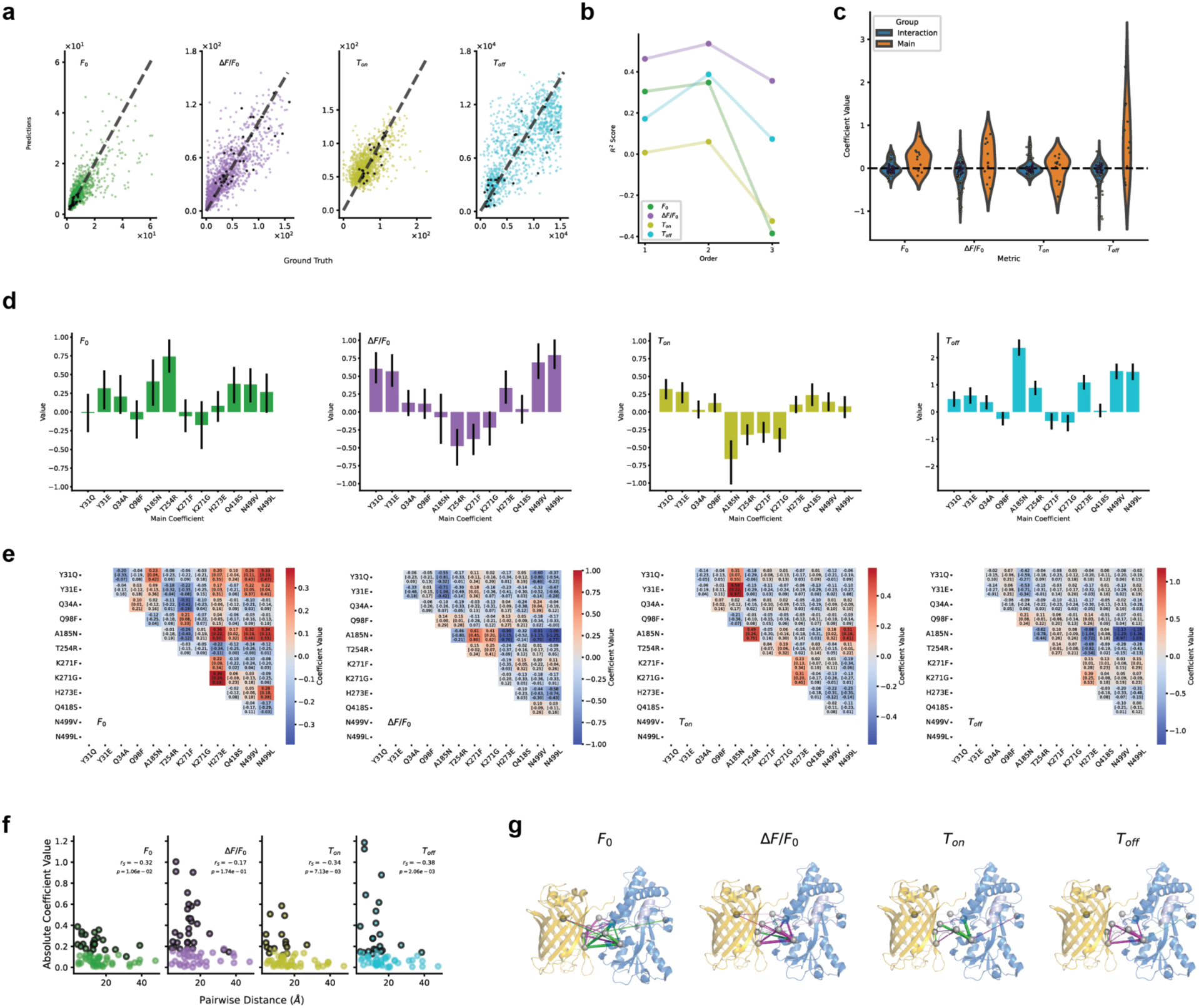
Analysis of mutation effects and spatial interactions. **a**) Measured variant mean values and corresponding predictions of a second-order GLM for each of the four response variables F0,(baseline), ΔF/F_0_ (fractional response), T_on_ (rise time constant), and T_off_ (decay time constant) (with the largest 1% of outliers clipped). The points in black correspond to variants that contain only a single mutation from the 12 mutations in the combinatorial screen. **b)** Cross-validated variance explained for GLMs of different orders: 1 (main effects only), 2 (main effects and all pairwise interactions), 3 (main effect and all pair and triplet interactions). The order 2 model was used in this study. The yellow (in main) and blue (in interaction) points represent statistically significant coefficients. **c)** Distributions of coefficients for main effects (orange) and interactions (blue) for the response variables. 34 of 48 main effects and 92 of 252 interactions were individually statistically significant at 95% confidence. **d)** Main effects of each mutation and bootstrapped 95% confidence intervals. **e)** Interaction terms for each mutation pair and bootstrapped 95% confidence intervals. **f)** Pairwise distances and coefficient magnitudes for interaction terms, and corresponding Spearman’s correlation coefficients. The outlined points represent statistically significant coefficients. **g)** iGluSnFR3 glutamate-bound crystal structure solved in this work (PDB: 9FBU), with significant interactions overlaid for each response variable. Green links denote positive interactions. Purple links denote negative interactions. Line thickness proportional to coefficient magnitude. On average across all metrics, the number of statistically significant interactions between residues are of the following types: 3%/8% FP-FP, 8%/32% FP-BP, 6%/14% BP-BP, 12%/24% BP-Linker, and 1%/3% Linker-Linker (where the numerator is the percentage of statistically significant interactions of the total number of interaction coefficients, and denominator is the percentage of possible interactions of the given interaction type of the total interactions).

**Supplementary Figure 2:**
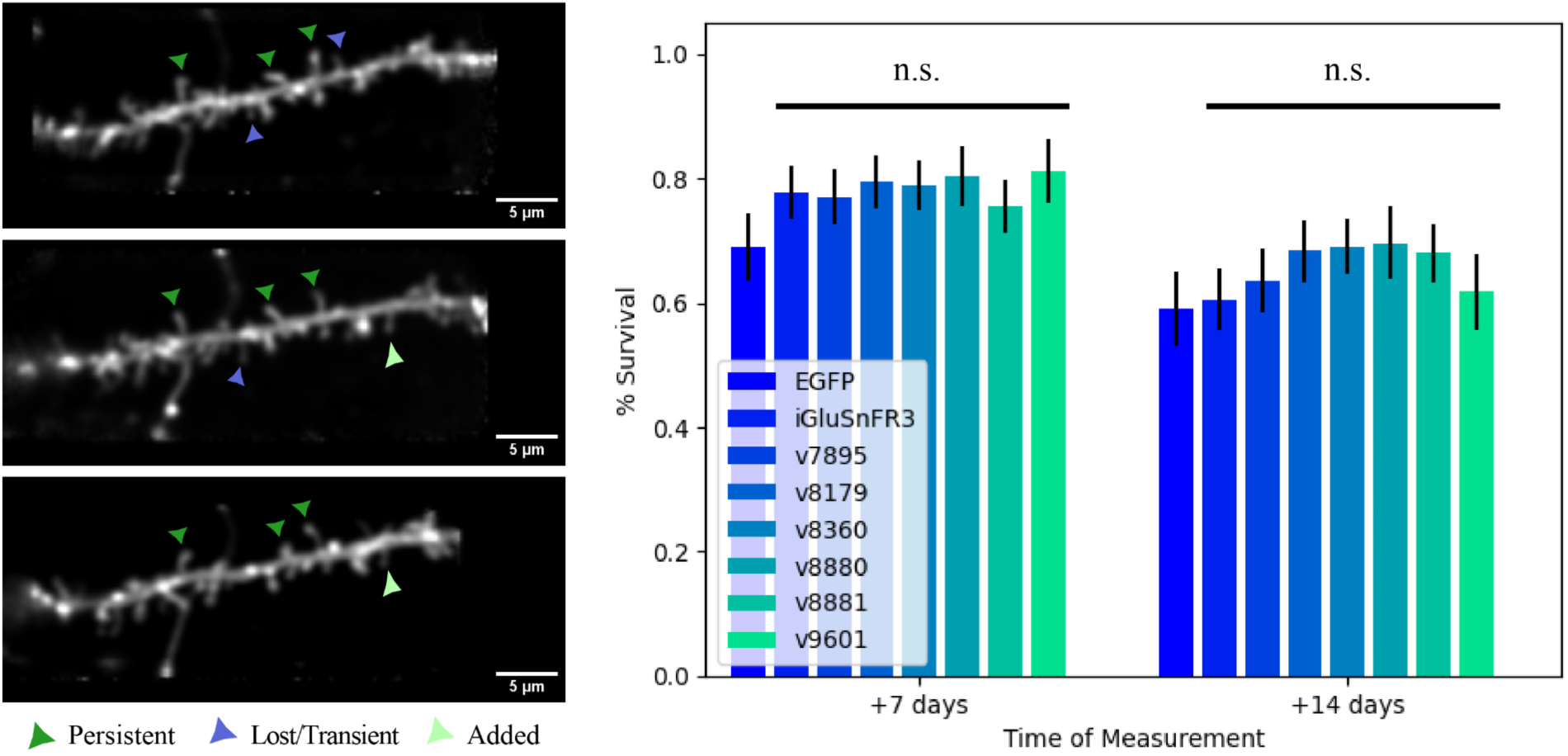
Spine turnover of iGluSnFR variants. Left, example measurements of spine turnover over 3 measurement days at 1-week intervals, for a v8880-expressing neuron. Right, Surviving fraction of spines over 3 measurement days at 1-week intervals for the iGluSnFR variants screened in vivo, and membrane-tagged EGFP control. Mean +/-Binomial standard error. N= [246 EGFP; 257 iGluSnFR3; 260 v7895; 163 v8179; 221 v8360; 178 v8880; 101 v8881; 142 v9601] spines at initial timepoint. A log-rank test (Kaplan-Meier model) was used to test total survival curves of each variant against EGFP. Chi-squared tests were performed to test the individual measurements taken at +7 days and +14 days against corresponding EGFP measurements. All p-values >0.05.

**Supplementary Figure 3:**
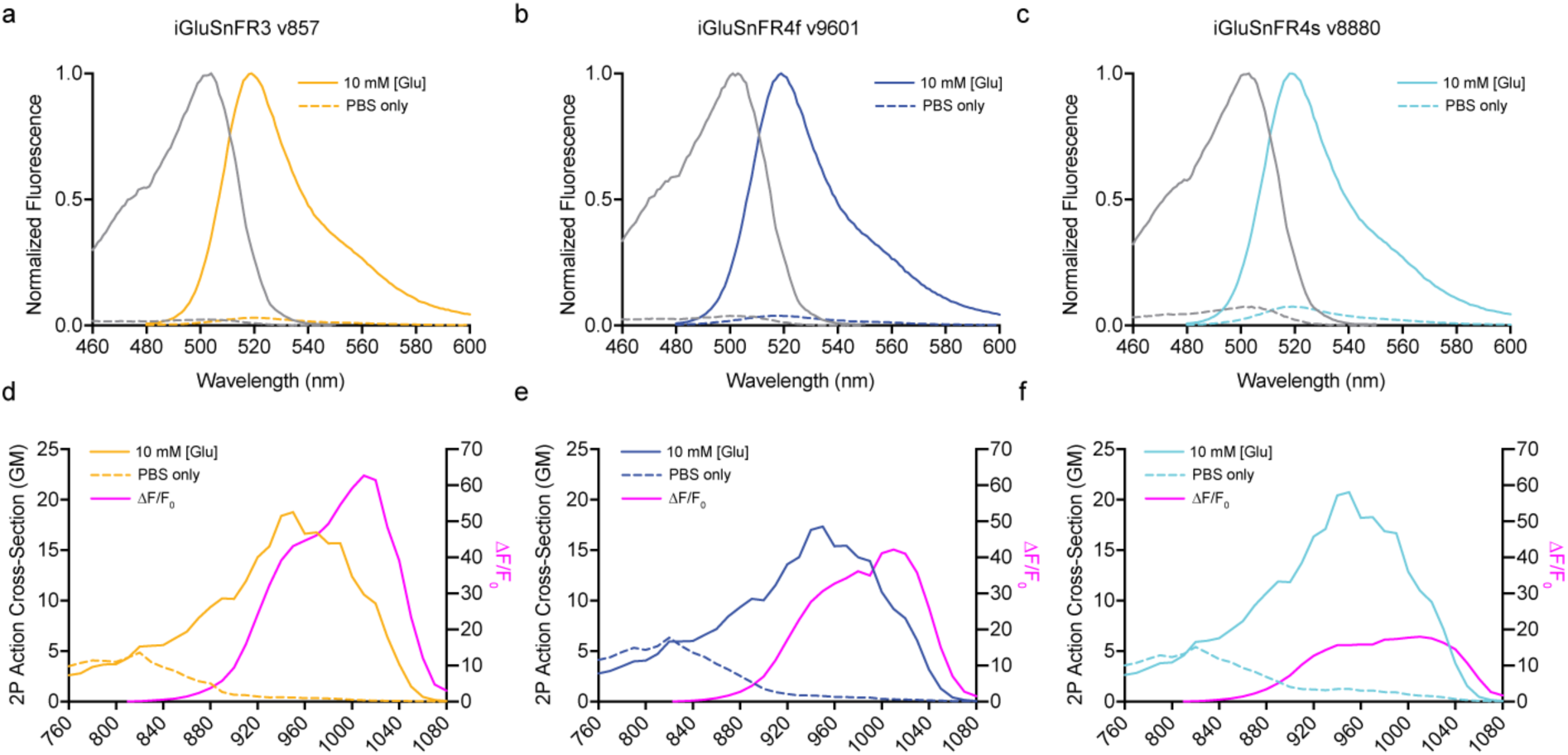
1P and 2P spectra of iGluSnFR variants. **a-c**) One-photon excitation and emission spectra of iGluSnFR3 v857, iGluSnFR4f, and iGluSnFR4 in the presence (10 mM) and absence of glutamate. **d-f)** Two-photon spectra and ΔF/F_0_ of the iGluSnFR4 variants in the presence (10 mM) and absence of glutamate.

**Supplementary Figure 4:**
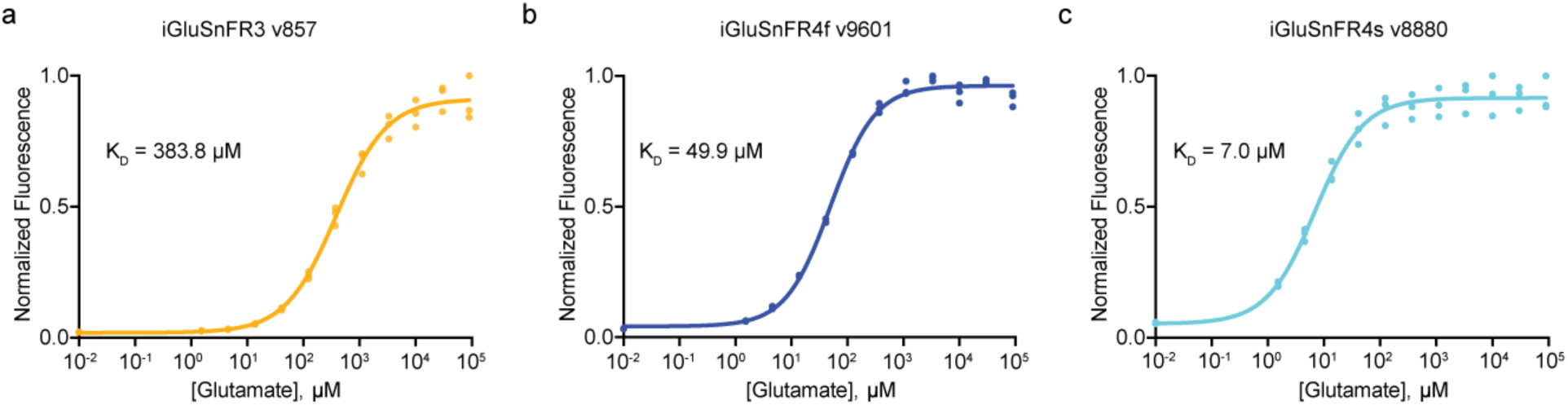
Glutamate titration of purified soluble proteins. **a**) iGluSnFR3.v857 (left), **b)** iGluSnFR4f (middle) and **c)** iGluSnFR4s (right) (pH 7.3 for all), with corresponding fits and dissociation constants (K_D_). N = 3 titration series of a single protein sample for each variant.

**Supplementary Figure 5:**
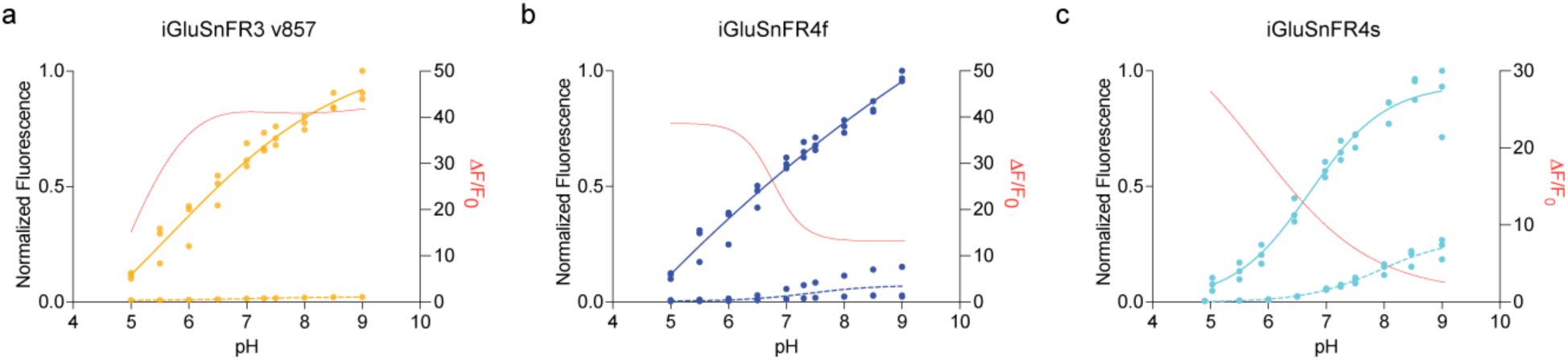
pH titration of purified soluble proteins. **a**) iGluSnFR3.v857 (left), **b)** iGluSnFR4f (middle) and **c)** iGluSnFR4s (right). Solid lines: saturating glutamate (10 mM, pH 7.3 buffered in PBS); Dotted lines: absence of glutamate. Sigmoidal fits are overlaid. Red lines show ΔF/F_0_ as a function of pH. All measurements were made using purified soluble protein; N = 3 titration series of a single protein sample for each measurement.

**Supplementary Figure 6:**
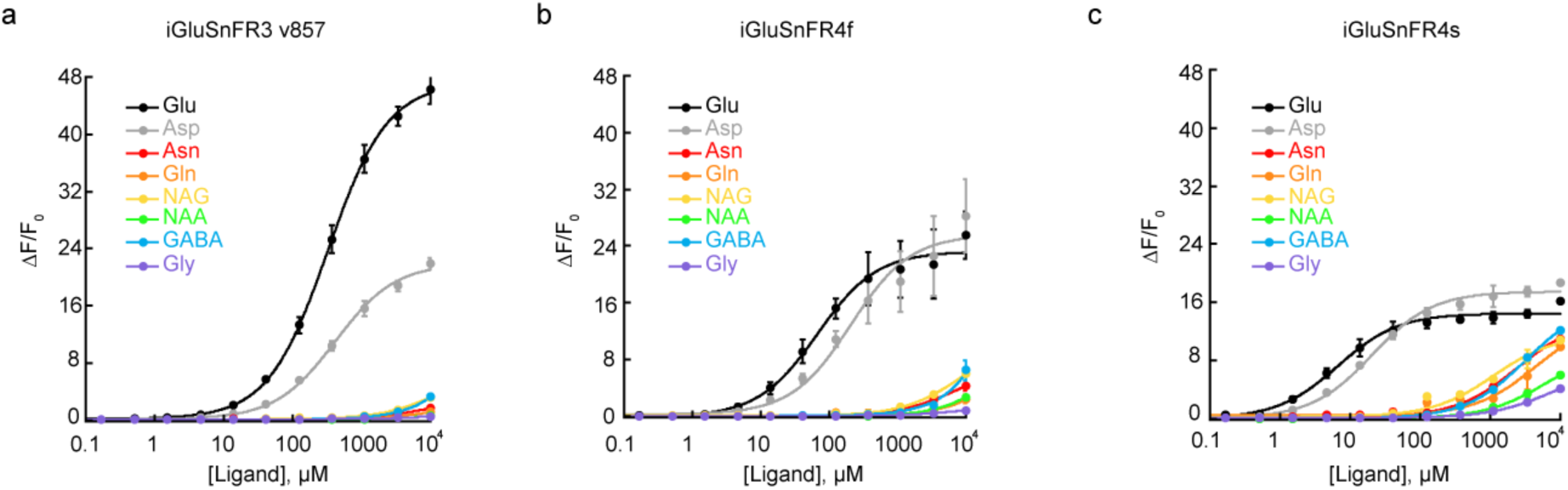
*In vitro* specificity of soluble proteins. ΔF/F_0_ of iGluSnFR3 **(a)**, iGluSnFR4f **(b)** and iGluSnFR4s **(c)** for titrations of selected L-amino acids, neurotransmitters, and other drugs (pH 7.3, buffered in PBS). All measurements were made using purified soluble protein; N = 3 titration series of a single protein sample for each measurement.

**Supplementary Figure 7:**
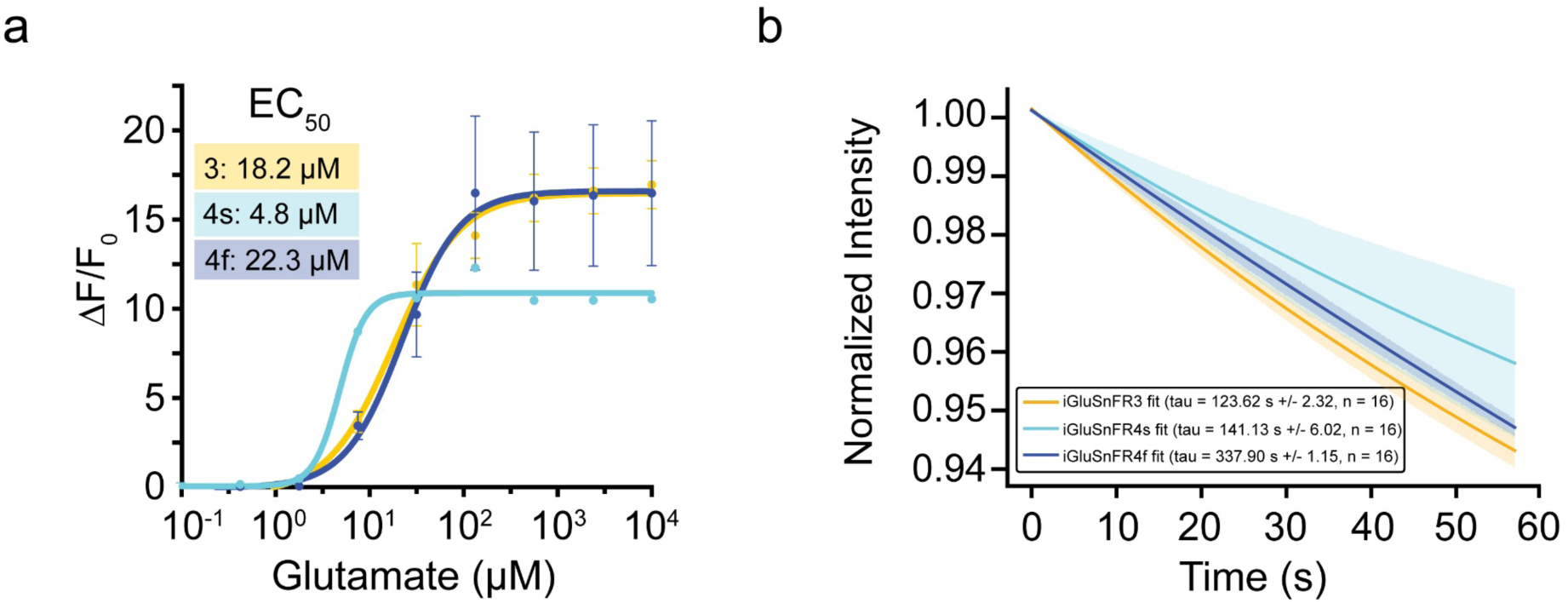
Glutamate titrations and photobleaching of iGluSnFR4 variants on the surface of cultured neurons. **a**) Titration curves for iGluSnFR4 variants expressed on the surface of cultured neurons. Data points represent the mean of N = 3 replicates, with error bars indicating SEM. A sigmoidal curve was fitted to the data. **b)** Photobleaching profiles of the three biosensors on the surface of cultured neurons under 10 Hz stimulation with 100 ms exposure per frame (570 frames over 57 seconds). The data were fitted using an exponential decay model. N = 16 replicates, with the shaded region representing SEM.

**Supplementary Figure 8:**
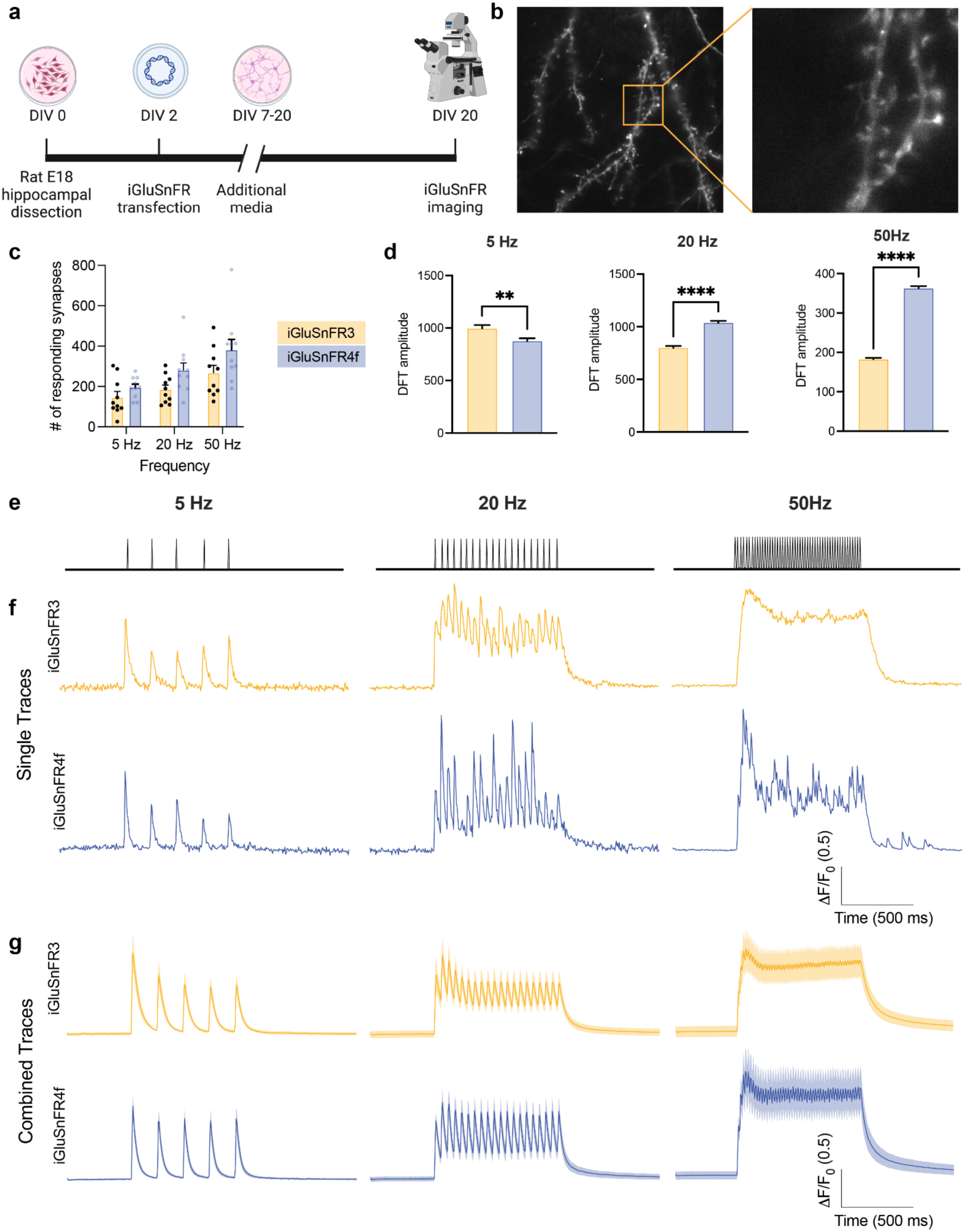
High frequency stimulation in hippocampal neuron culture. **a**) Experiment scheme. Hippocampi from E18 rat pups were dissected (DIV 0) and transfected with iGluSnFR plasmids at DIV 2. Neurons were given additional media from DIV 7-20, then iGluSnFR imaging was performed on DIV 20. Made with Biorender. **b)** Representative field of view of a dendritic arbor expressing iGluSnFR. **c)** Number of synapses responding to either 5 Hz, 20 Hz, or 50 Hz stimuli for iGluSnFR3 and iGluSnFR4f. n=10 neurons for each condition. Data shown is the mean±standard error of the mean. **d)** Power spectral amplitude of evoked responses at 5 Hz, 20 Hz and 50 Hz. Bar graphs represent the mean±standard error for n=10 neurons per variant. Unpaired t-test; **p<0.01, ****p<0.0001. **e)** Representative stimulation paradigms for 5 Hz/5 stimuli, 20 Hz/20 stimuli, and 50 Hz/50 stimuli. **f)** Traces from single synapses for both iGluSnFR3 and iGluSnFR4f in response to either 5 Hz, 20 Hz or 50 Hz stimuli. **g)** Combined traces from n=10 neurons for iGluSnFR3 and iGluSnFR4f in response to either 5 Hz, 20 Hz or 50 Hz stimuli. Data shown is the mean±standard deviation.

**Supplementary Figure 9.**
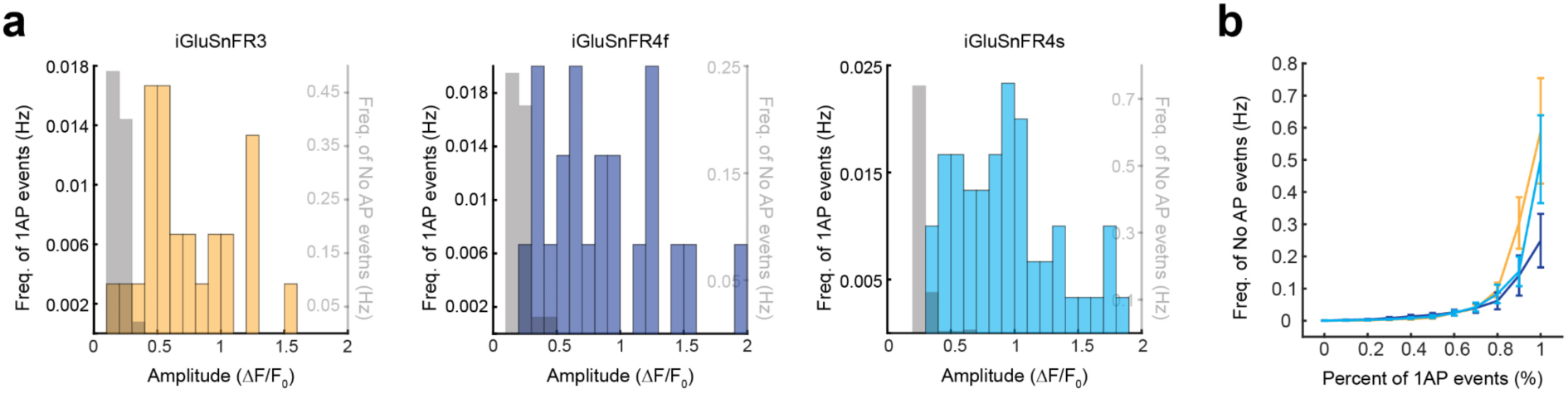
Detection of single action potentials by iGluSnFR variants. **a**) Amplitude histograms for single-AP events in V1 axons for iGluSnFR3, 4f, and 4s, and events without associated APs at the corresponding amplitude thresholds in grey. **b)** Frequency of false positives (no AP events) as a function of fraction of true positive 1 AP events detected, for iGLuSnFR3 (yellow), 4f (navy blue) and 4s (cyan).

**Supplementary Figure 10:**
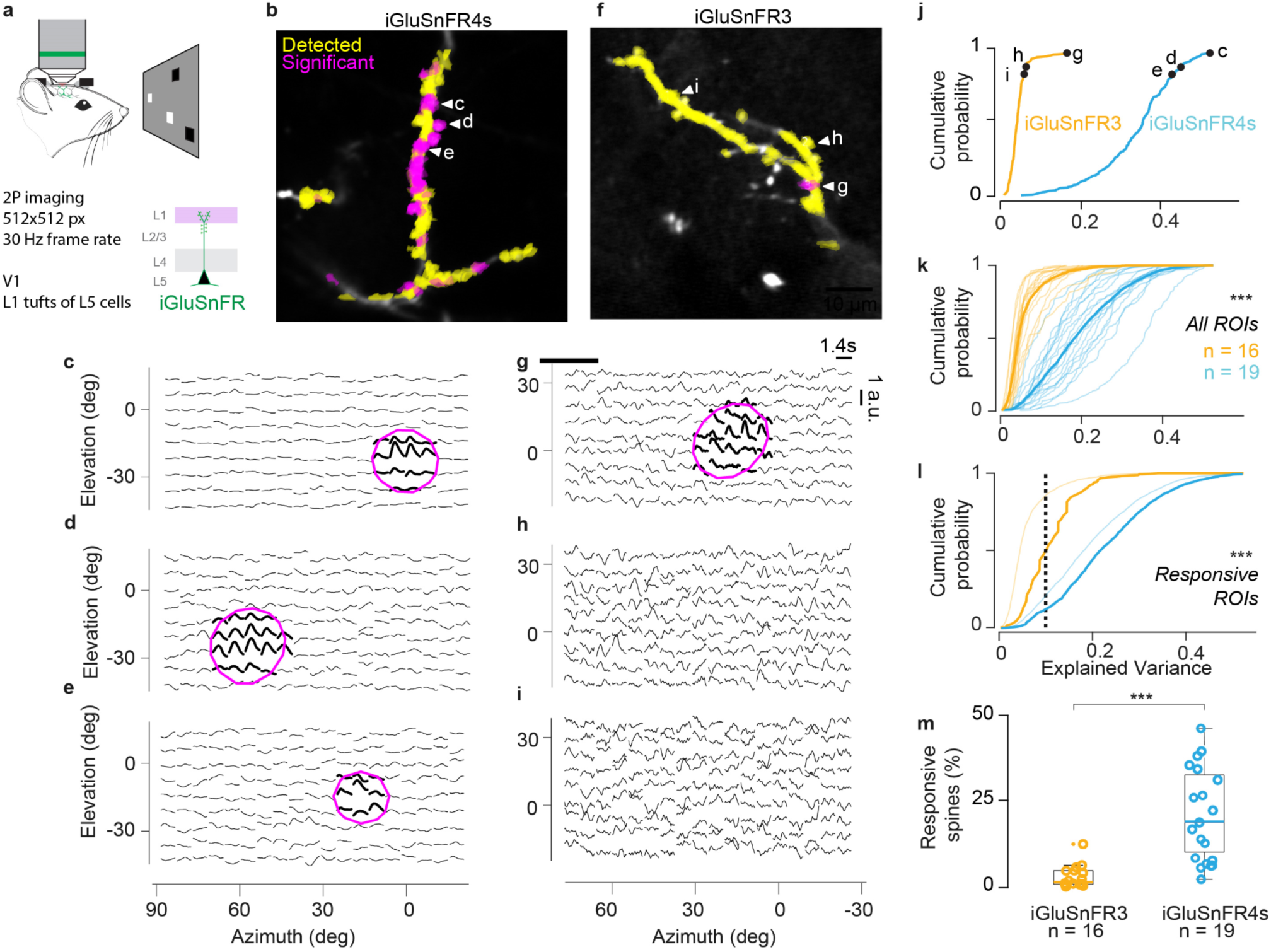
Video-rate imaging in visual cortex. **a**) Mice passively viewed sparse noise stimuli while imaging apical dendrites of L5 neurons in mouse V1 sparsely labeled with iGluSnFR3 or 4s. **b)** Example mean image of an apical dendrite expressing iGluSnFR4s. Active ROIs with significant receptive fields (magenta; p<0.005 vs circular shift null and >10% variance explained by stRF) and all others detected by Suite2p (yellow) are overlaid to the normalised mean fluorescence image. **c–e)** Spatiotemporal receptive fields (stRFs) fit for ROIs labelled in (b), normalized to peak response. Guassian fit overlaid(magenta ellipse and thick traces). **f-i)** Same as (b-e) for example dendrite expressing iGluSnFR3. **j)** Cumulative probability distributions of the explained variance from all the ROIs detected on the two example dendrites shown in b and f. The example RF shown in c-e and g-i were randomly selected as those at the 85th, 90th and 100th percentile of the distribution (***p < 0.001, Kolmogorov-Smirnov test). **k)** Cumulative distribution of RF explained variance from all recorded dendrites (thin lines) and the average across dendrites (thick lines) for iGluSnFR3 (orange, 16 dendrites), and iGluSnFR4s (cyan, 19 dendrites). (***p < 0.001, Kolmogorov-Smirnov test). **l)** Average cumulative probability distributions of the explained variance of all stRF (thin lines, same as k) or of stRF explaining more variance than chance ROIs only (thick line), for iGluSnFR3 (orange) and iGluSnFR4s (cyan). The dotted line shows the threshold of explained variance fraction (0.1) used to determine visual responsiveness. (***p < 0.001, Kolmogorov-Smirnov test). **m)** The percentage of responsive ROIs was significantly higher for dendrites expressing iGluSnFR4s (cyan) compared to iGluSnFR3 (orange, ***p < 0.001, linear mixed-effects model). The iGluSnFR3 population consisted of 16 dendrites, pooled from 7 neurons in a total of 3 mice. The iGluSnFR4s population consisted of 19 dendrites, pooled from 3 neurons in a total of 2 mice.

**Supplementary Figure 11:**
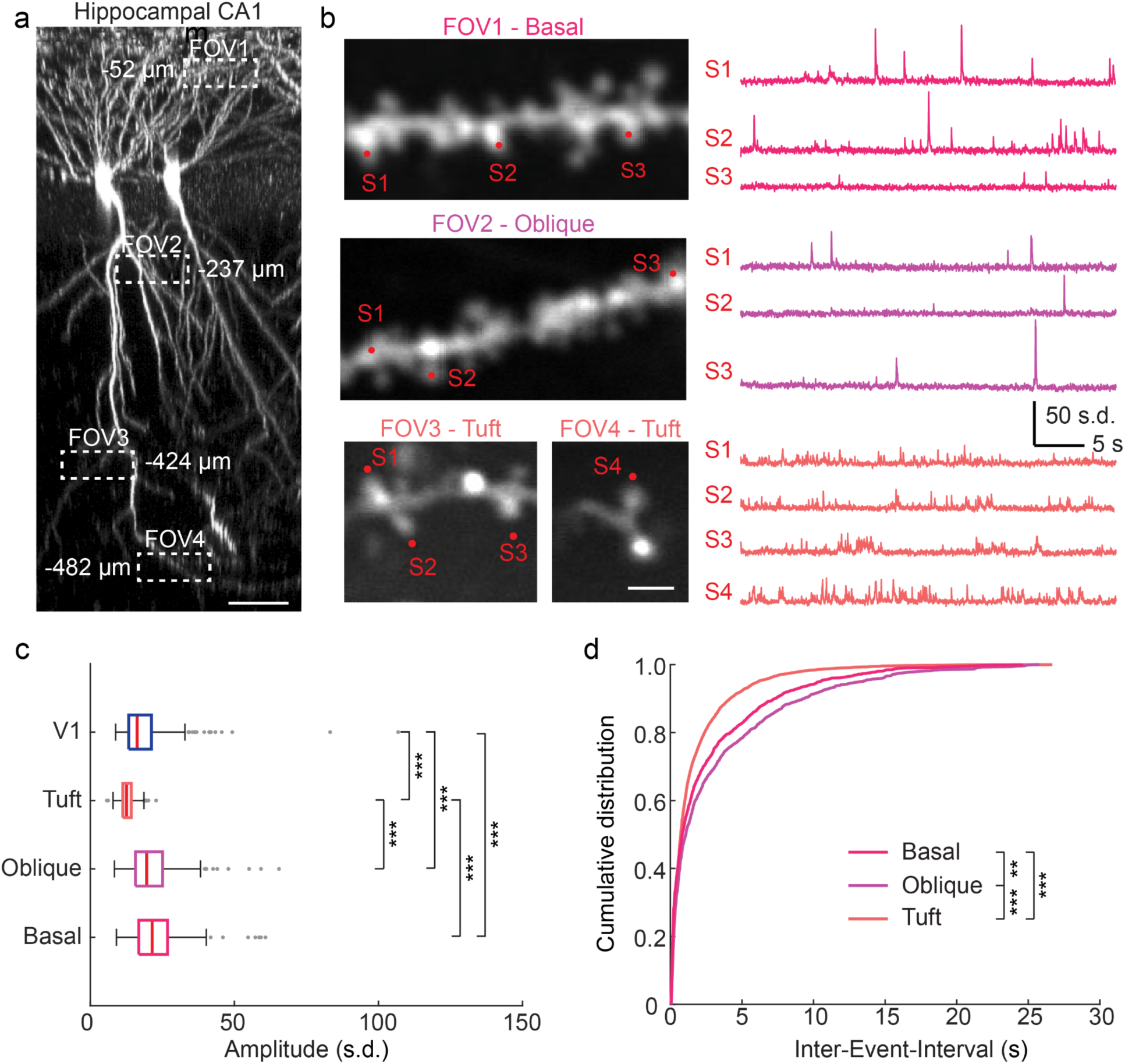
iGluSnFR4f imaging in hippocampus CA1. **a**) Z-projection of iGluSnFR4f labeled CA1 pyramidal neurons and locations of imaged dendritic segments. The number next to the box indicates the depth below the coverslip. Scale bar, 50 μm.**b)** Two-photon images of dendritic segments from basal, oblique, and tuft dendrites of CA1 pyramidal neurons (left) and z-scored iGluSnFR4f traces from denoted spines (right). Scale bar, 2 μm. **c)** Amplitudes of synaptic glutamate signals from basal (n = 265 ROIs from 4 cells), oblique (n = 286 ROIs from 4 cells), and tuft dendrites (n = 444 ROIs from 4 cells) of pyramidal neurons in CA1 and L2/3 dendrites (n = 465 ROIs from 4 cells) in V1. (***P < 0.001; Kruskal-Wallis test with Dunn’s test for multiple comparisons) **d)** Distribution of Inter-Event-Interval for glutamate signals from different dendritic domains of CA1 pyramidal neurons (**P < 0.01; ***P < 0.001; Kolmogorov–Smirnov test).

**Supplementary Table 1:** Photophysical characterization of iGluSnFR3, iGluSnFR4f, and iGluSnFR4s.

| Variant Name | iGluSnFR3 |  | iGluSnFR4f |  | iGluSnFR4s |  |
| --- | --- | --- | --- | --- | --- | --- |
|  | APO | SAT | APO | SAT | APO | SAT |
| 1-photon Excitation maxima $\lambda_{\text{exc}}$ (nm) | 502 | | 502 | | 502 | |
| 1-photon Emission maxima $\lambda_{\text{exc}}$ (nm) | 522 | | 522 | | 522 | |
| 1-photon $\Delta F/F$ | 47.1 | | 22.9 | | 17.2 | |
| $K_d$ ($\mu\text{M}$) [95% C.I.] | 383.8 $\mu\text{M}$ [325.2, 453.0] | | 49.9 $\mu\text{M}$ [44.5, 56.1] | | 7.0 $\mu\text{M}$ [5.6, 8.7] | |
| Apparent Hill coefficient ($n_H$) [95% C.I.] | 0.9 [0.8, 1.1] | | 1.1 [1.0, 1.2] | | 1.0 [0.8, 1.2] | |
| Apparent pKa | 7.0 | 5.7 | 7.4 | 6.5 | 7.9 | 6.7 |
| Extinction Coefficient $\epsilon$ ($\text{M}^{-1}\text{cm}^{-1}$) | 2055 | 35000 | 3510 | 32920 | 4430 | 40440 |
| Quantum Yield $\Phi$ | 0.29 | 0.88 | 0.27 | 0.89 | 0.46 | 0.88 |
| 1-photon Brightness (x1000) | 0.60 | 30.8 | 0.95 | 29.3 | 2.0 | 35.6 |
| 2-photon Brightness $F_2(\text{GM})$, (950 nm / 1030 nm) | 18.8 / 6.4 | | 17.3 / 6.1 | | 20.7 / 7.2 | |
| 2-photon $\Delta F/F$, (950 nm / 1030 nm) | 43.1 / 48.4 | | 30.6 / 35.7 | | 15.7 / 16.3 | |

**Supplementary Table 2:** Sequences of variants selected for *in vivo* testing.

| Variant name | Mutations (GltI-N cp-mVenus GltI-C) |
| --- | --- |
| iGluSnFR3 v857 | Template |
| iGluSnFR4s (v8880) | Y31Q Q98F K271F N499V |
| iGluSnFR4f (v9601) | Y31Q Q98F S182V K271F N499V |
| v7895 | Y31Q Q98F N499V |
| v8179 | Y31Q Q34A Q98F N499V |
| v8360 | Y31Q Q34A Q98F K271G Q418S N499V |
| v8376 | Y31E Q34A Q98F K271G N499V |
| v8881 | Y31Q A185N K271G N499V |

**Supplementary Table 3:**
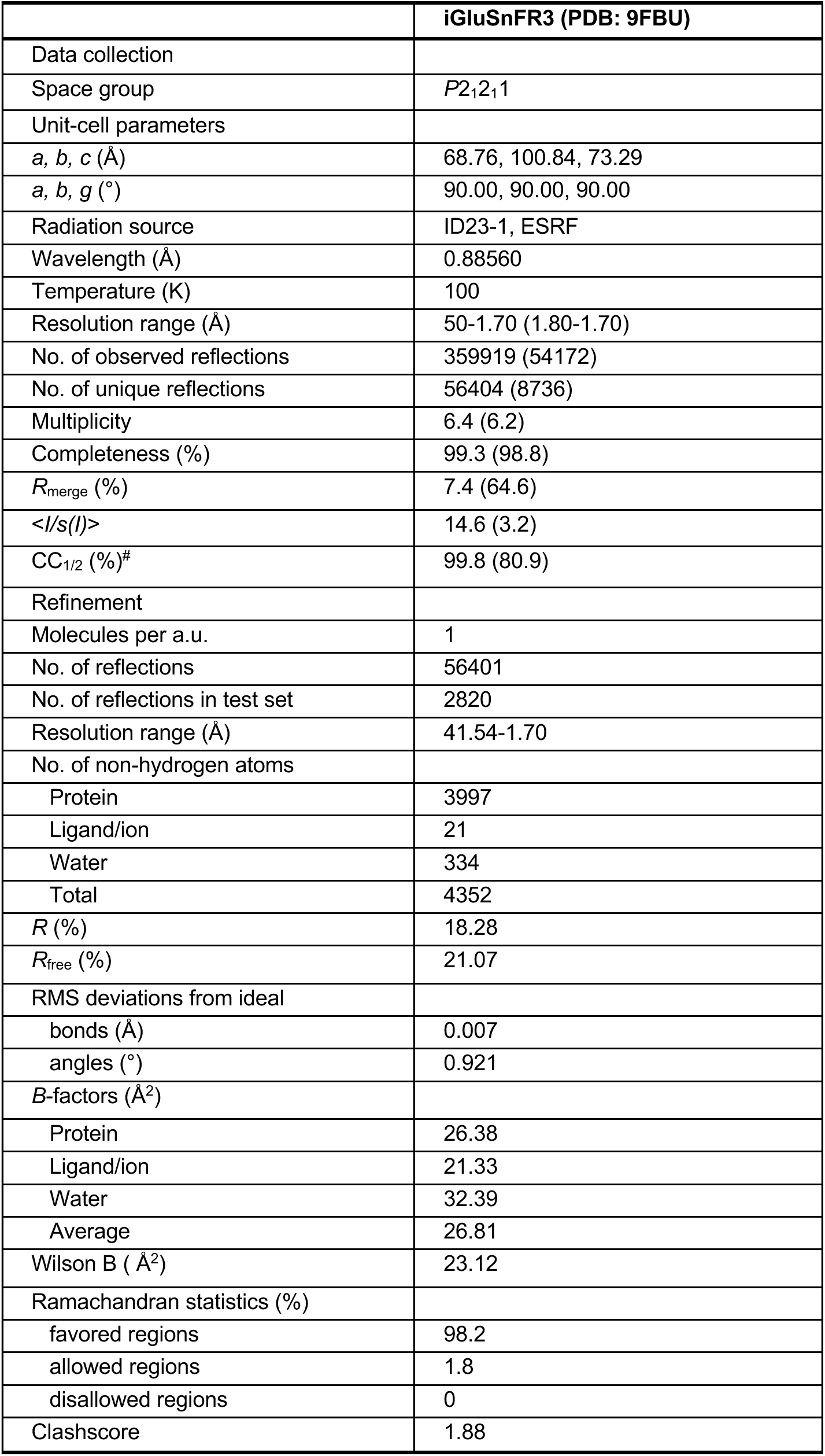
X-ray crystallography data collection and refinement statistics. Values in parentheses are for the highest resolution shell, as implemented in XDS.

|  | iGluSnFR3 (PDB: 9FBU) |
| --- | --- |
| Data collection |  |
| Space group | *P*2<sub>1</sub>2<sub>1</sub>1 |
| Unit-cell parameters |  |
| *a*, *b*, *c* (Å) | 68.76, 100.84, 73.29 |
| *a*, *b*, *g* (°) | 90.00, 90.00, 90.00 |
| Radiation source | ID23-1, ESRF |
| Wavelength (Å) | 0.88560 |
| Temperature (K) | 100 |
| Resolution range (Å) | 50-1.70 (1.80-1.70) |
| No. of observed reflections | 359919 (54172) |
| No. of unique reflections | 56404 (8736) |
| Multiplicity | 6.4 (6.2) |
| Completeness (%) | 99.3 (98.8) |
| *R*<sub>merge</sub> (%) | 7.4 (64.6) |
| <*I*/*s*(*I*)> | 14.6 (3.2) |
| CC<sub>1/2</sub> (%)<sup>#</sup> | 99.8 (80.9) |
| Refinement |  |
| Molecules per a.u. | 1 |
| No. of reflections | 56401 |
| No. of reflections in test set | 2820 |
| Resolution range (Å) | 41.54-1.70 |
| No. of non-hydrogen atoms |  |
| Protein | 3997 |
| Ligand/ion | 21 |
| Water | 334 |
| Total | 4352 |
| *R* (%) | 18.28 |
| *R*<sub>free</sub> (%) | 21.07 |
| RMS deviations from ideal |  |
| bonds (Å) | 0.007 |
| angles (°) | 0.921 |
| *B*-factors (Å<sup>2</sup>) |  |
| Protein | 26.38 |
| Ligand/ion | 21.33 |
| Water | 32.39 |
| Average | 26.81 |
| Wilson B ( Å<sup>2</sup>) | 23.12 |
| Ramachandran statistics (%) |  |
| favored regions | 98.2 |
| allowed regions | 1.8 |
| disallowed regions | 0 |
| Clashscore | 1.88 |

Supplementary Video 1: Example optical mini recording for iGluSnFR3, iGluSnFR4f, and iGluSnFR4s. Cyan denotes structure, and is the raw recording low pass filtered at 4 Hz, then gamma corrected (0.5). The residual (i.e. high-pass filtered recording) is shown in red (no gamma correction). Look up table for both channels are matched across the three videos. Video is played at 0.5x of the original recordings.

Supplementary Video 2: Average 2P-AO recordings of evoked thalamocortical bouton activity (mean of 60 one-second intervals) in vS1 Layer 4 at 5 Hz stimulus frequency.

## Methods

### Data availability

The data and code used to produce the figures are included in the code and data supplement (in preparation).

### Reagent availability

Bacterial (pRSET) and mammalian (pAAV) expression vectors, including different promoters, Cre- and Flp- dependent vectors, and both the NGR and PDGFR membrane targeting sequences, are available from Addgene (IDs: 234435-234454).

### Competing Interests Statement

The authors have no competing interests related to this work.

### Statistics and reproducibility

Values and errors reported through the text (“### ± ###”) are mean ± SEM unless otherwise noted. Statistical analyses were performed in Graphpad Prism 8, Python, and Matlab, as described in the text. Data and code used to produce the figures, and instructions for generating them, are provided in the data supplement (in preparation).

### Animal care and use statement

All experimental procedures involving animals were performed in accordance with protocols approved by the Institutional Animal Care and Use Committee at the respective institute (HHMI Janelia Research Campus, Allen Institute, University of California San Diego, TUM, University College London, St. Jude Children’s Research Hospital). All procedures performed in the United States conform to the National Institutes of Health (NIH) Guide for the Care and Use of Laboratory Animals. All procedures performed at TUM were approved by the state government of Bavaria, Germany. All procedures performed at UCL conform to the Animals Scientific Procedures Act (1986), and were performed under personal and project licenses released by the Home Office following appropriate ethics review.

*HHMI Janelia Research Campus: protocols 19-176, 22-0214.01*

*Allen Institute: Protocol 2109*

*University of California, San Diego: S02174M*

*Technical University of Munich:* 2532.Vet_02-23-24 and 2532.Vet_02-21-121

*University College London: Animal License PP3929312*

*St. Jude Children’s Research Hospital:* Protocol no. 3193

This research has complied with all applicable ethical regulations.

### Library Generation

We first performed random saturating mutagenesis at 41 targeted amino acid residues within iGluSnFR3, screening 96 random variants per site and sequencing only variants that excelled along one or more response variables. The mammalian expression vector pAAV.hSyn.iGluSnFR3.v857.GPI (Addgene #178331) and a related variant, iGluSnFR3.v867.GPI, were used as templates for site-directed mutagenesis. This vector contains a BglII restriction site at the N-terminus and a PstI restriction site at the C-terminus, flanking the iGluSnFR3 gene. To introduce mutations, overlapping internal degenerate-codon primers were designed for each site-specific change. Mutagenesis was performed using overlapping paired amplicons, which were assembled into the BglII and PstI digested pAAV.hSyn vector using a commercial HiFi DNA Assembly Mix. The assembled reaction products were transformed into NEB STABL2 chemically competent E. coli cells (New England Biolabs). Transformants were plated on LB agar plates supplemented with ampicillin (100 mg/L) and incubated overnight at 37°C. For each mutagenesis site, 96 colonies were picked and inoculated into 2.6 mL of 2x-YT media containing ampicillin (100 mg/L). The cultures were grown for 24 hours at 37°C with shaking at 225 rpm. Bacterial cells were then pelleted by centrifugation, and plasmid DNA was isolated using a miniprep kit (Qiagen). The DNA concentrations were normalized to 60 ng/µL based on absorbance readings at 260 nm using a Tecan Infinite M1000Pro microplate reader. Normalization and dilution of plasmid DNA were automated using a Hamilton Nimbus liquid handling system. Plasmids were electroporated into neuronal cell cultures along with unmutated controls. Variants with largest positive effect in each plate were selected for Sanger sequencing (Genewiz). Amino acid mutations that appeared two or more times in a plate were selected for further screening.

Top-performing single-site mutations (Y31Q, Y31E, Q34A, Q98F, A185N, T254R, K271F, K271G, H273E, Q418S, N499L, N499V) were selected based on their high performance on at least one of the four response metrics. A total of 12 specific residue mutations at 9 positions were chosen for combinatorial testing, encompassing all possible combinations (1728). Although the initial saturation screen was performed on iGluSnFR3.v857 and iGluSnFR3.v867 in parallel, due to poor photostability observed in all derivatives of iGluSnFR3.v867, combinatorial mutagenesis was performed using only pAAV.hSyn.iGluSnFR3.v857.GPI as the template. To generate these combinations, two overlapping amplicons spanning the iGluSnFR3 gene were amplified via PCR. The PCR products were then purified and ligated into the pAAV.hSyn vector that had been digested with BglII and PstI restriction enzymes. The assembled constructs were transformed into NEB STABL2 chemically competent E. coli cells (New England Biolabs), resulting in the isolation of 1728 unique isothermal assembly products in 96-well cultures. These bacterial cultures were grown (225 rpm, 37°C), pelted, and miniprepped. The extracted plasmids were confirmed by Sanger sequencing to verify the presence of the intended combinations. Following verification via Sanger sequencing, the combinatorial plasmids were normalized to a concentration of 60 ng/µL and arrayed for subsequent electroporation into neuronal cell cultures.

A total of 3365 variants were screened. 1640 unique variants were screened in the saturation library (41 sites * 20 residues * 2 templates, assuming full residue coverage by the 96 random samples per variant), 1728 in the combinatorial library (of which 13 were already present in the saturation screen), and 10 candidate fast variants v8880+(S70A, S70T, S182L, S182V, or Y209F) and v8376+(S70A, S70T, S182L, S182V, or Y209F).

### Neuron culture screen

Neonatal rat pups (Charles River Laboratory) were euthanized, and the hippocampal/cortical cell culture was obtained. Tissue dissociation was carried out using papain (Worthington) in 10 mM HEPES (pH 7.4) prepared in Hanks’ Balanced Salt Solution, incubating at 37°C for 30 minutes. The resulting cell suspensions were triturated with a Pasteur pipette and filtered through a 40-µm strainer. For transfection, 5x10^5 viable cells were mixed with 400 ng of plasmid DNA and nucleofection solution in a 25 µL electroporation cuvette (Lonza) and electroporated following the manufacturer’s instructions. For the field stimulation assay, neurons were plated at a density of 1x10^5 cells per well in poly-D-lysine (PDL) coated, 96-well glass bottom plates (MatTek, #1.5 cover glass). Cells were plated in 100 µL of a medium composed of a 4:1 ratio of NbActiv4 (BrainBits) and plating medium (28 mM glucose, 2.4 mM NaHCO3, 100 µg/mL transferrin, 25 µg/mL insulin, 2 mM L-glutamine, 100 U/mL penicillin, 10 µg/mL streptomycin, and 10% FBS in MEM). On the following day, 190 µL of NbActiv4 medium was added to each well, and plates were incubated at 37°C with 5% CO_2_ for 12-15 days before imaging. Typically, 8 wells of a 96-well plate were used to electroporate iGluSnFR3.v857 as a control, while the remaining wells (4 per construct) were used for the constructs of interest. To minimize edge effects, the first and last columns of the plate were not utilized. Each well was imaged under 1 AP and 20 AP field stimulation, as previously described^26^. We calculated scalar performance metrics per well, including brightness, photostability, number of responsive pixels, ΔF/F_0_ relative to a 1s-window before stimulation, and rise and decay rates fit to the mean pixel traces. Metrics for each well were normalized to the in-plate iGluSnFR3.v857 controls for analysis. Summary data and code used for analysis are included in the data supplement.

### Modeling of mutation effects

Analysis of the combinatorial field stimulation screening data was done using a Generalized Linear Model (GLM) approach implemented using the *statsmodels*^48^ *(v0.14.2)* package in Python 3.12.2. The input to the GLM is a 9-length vector representation of a protein variant in a categorical format - there are 9 categories (covering the total 9 sites) and each site can either have no mutation, mutation 1 or mutation 2 for some sites. This covers the total of 12 mutations with 3 of the sites having 2 possible mutations, and the other 6 having 1 possible mutation. The labels provided to the GLM are the values of the sensor performance metrics: ΔF/F_0_, F_0_, T_off_ or T_on_ - corresponding to a given variant. For each variant multiple measurements (corresponding to imaging on different days, screens, plates, or wells) were used and no averaging was done. Model parameters for 20 AP field stimulus data is shown in Fig S1; 1 AP fits exhibit lower SNR but were otherwise qualitatively similar. Negative values in ΔF/F_0_ were clipped to 0. The appropriate link function and exponential dispersion model (EDM) family was chosen using an exhaustive search and the ones with the best fit (as assessed by smallest RMSD on predictions) for each metric were selected. The Poisson family with the log link function was chosen for F_0_, and ΔF/F_0_, and the Gaussian family with the log link function was chosen for T_on_ and T_off_.

Bootstrapping was used to generate confidence intervals of the coefficients in the GLM fits. The fits were repeated by randomly selecting N variants (with repetition) over the dataset, where N is the total size of the dataset. 95% confidence intervals for each coefficient were estimated from the resulting distributions. Coefficients were marked as statistically significant if 0 does not lie within their 95% confidence intervals.

Distances between residues were calculated using the ‘get_distanc’ function from PyMol which calculates the Euclidean distance between the coordinates of two atoms. For the pairwise distance between mutations this was calculated between the closest atom pair from the residues at those sites.

### Protein crystallization, X-ray diffraction, and structure determination

Soluble iGluSnFR3 (pRSET iGluSnFR3.v857, Addgene # 175186) was subcloned, expressed and purified as previously described for AspSnFR^49^. Crystallization was performed at 20°C using the vapor-diffusion method. The protein at a concentration of 18.0 mg/ml in 50 mM HEPES pH 7.3, 50 mM sodium chloride was supplemented with 90 mM L-glutamate prior to crystallization. Crystals of iGluSnFR3 in complex with L-glutamate were grown by mixing equal volumes of protein solution and precipitant solution containing 26% (m/v) PEG 1500. Before flash-cooling in liquid nitrogen the crystals were briefly washed in a cryoprotectant solution consisting of the reservoir solution supplemented with 20% (v/v) glycerol.

Single crystal X-ray diffraction data were collected at 100 K on the ID23-1 beamline at the ESRF (Grenoble, France). All data were processed with XDS^50^. The structure of iGluSnFR3 was determined by molecular replacement (MR) with Phaser^51^ using SF-iGluSnFR-S72A coordinates (PDB: 8OVO) as a search model. The final model was optimized in iterative cycles of manual rebuilding using Coot^52^ and refinement using Refmac5^53^ and phenix.refine^54^. Data collection and refinement statistics are summarized in Supplementary Table 3. Model quality was validated with MolProbity^55^ as implemented in PHENIX. Atomic coordinates and structure factors have been deposited in the Protein Data Bank under accession code 9FBU.

### Imaging optical minis

For imaging spontaneous release, 2x105 cells were plated onto PDL-coated, 35-mm glass bottom dishes (Mattek, #0 cover glass) in 120 mL of a 1:1 mixture of NbActiv4 and plating medium in the center of the plate. The next day, 2 mL of NbActiv4 medium was added to each plate. 50% of the medium was replaced with fresh medium at 4 and 7 DIV. Imaging was performed at 14 DIV. Prior to experiment, culture media was replaced with imaging buffer containing the following (in mM): 145 NaCl, 2.5 KCl, 10 glucose, 10 HEPES, pH 7.4, 2 CaCl2, 1 MgCl2, 100 mM sucrose (to enhance glutamate release) and 2 mM TTX (to block AP-evoked release). Images were captured with an inverted fluorescence microscope (Zeiss AXIO Observer 7) equipped with SPECTRA X light engine (Lumencore), a 63X oil objective (NA = 1.4, Zeiss), and a scientific CMOS camera (Hamamatsu ORCA-Flash 4.0). A FITC filter set (475/50 nm (excitation), 540/50 nm (emission), and a 506LP dichroic mirror (FITC-5050A-000; Semrock)) was used for all iGluSnFR variants in this study. For each neuron culture plate, 6-10 FOVs were chosen and imaged with constant light intensity at 12 mW/mm^2^. For each FOV, 3000 images (512x512 pixels) were acquired with Hamamatsu image acquisition software (HCImageLive) at 100 Hz. Signal (miniature glutamate release evoked fluorescence increase) was analyzed with a custom MATLAB script.

### Analysis of optical minis

Optical mini recordings were processed using the same Matlab script as in previous work^26^. SNR for each detected site was calculated as the amplitude of the third largest peak in the 30-second recording, divided by a spectral estimate of the noise amplitude for each trace. The decay time at each site was calculated by isolating peaks in the site’s activity trace and fitting an exponential decay function over the mean aggregated data over a time window following the peak. The median of each statistic across detected sites was calculated for each well, and the mean and SEM of the well medians is shown in Fig. 2d.

The pixelwise statistic used to generate the activity images (red channel in Figs 2a,e) is the skewness of the highpass filtered image normalized by the square root of the mean intensity. For data derived from a Poisson process (*e.g.* photon shot noise), the expected value of this statistic is 1; values above 1 reflect positive skewness deviations in excess of shot noise.

### Frequency response in hippocampal culture

Sprague Dawley rat hippocampal neurons were isolated at E18 (Envigo). Hippocampal tissue was dissected and maintained in chilled hibernate A media (Gibco; 1247501). This tissue was then incubated with 0.25% trypsin (Corning; 25-053 Cl) for 30 min at 37°C before trituration in DMEM (Gibco; 11965-118) supplemented with 10% fetal bovine serum (FBS, Thermofisher; 501527079) and penicillin-streptomycin (pen/strep, Thermofisher; MT-30-001-Cl). Dissociated neurons were plated on 18 mm glass coverslips (Warner instruments; 64-0734 [CS-18R17]) coated with poly-D-lysine (Thermofisher; ICN10269491). Neurons were initially plated in DMEM with 10% FBS and pen/strep for 2 hours at a density of 125,000 cells/well. Then, media was replaced with Neurobasal-A medium (Thermofisher; 10888-022) supplemented with 2% B-27 (Thermofisher; 17504001), 2mM Glutamax (Thermofisher; 35050031), and pen/strep. On DIV 2, neurons were transfected with 200ng of plasmid DNA encoding hSyn.iGluSnFR.NGR variants using lipofectamine LTX with PLUS reagent using 1ml of lipofectamine LTX and 0.5ml of PLUS reagent per well (Thermofisher; 15338100). Neurons were supplemented with media once a week and imaged at DIV 20.

Imaging was performed on an Olympus IX83 inverted microscope equipped with a X-cite XYLIS LED (Excelitas; XT720S), Olympus 100x/1.5 NA objective (UPLAPO100XOHR), and an ORCA Fusion BT sCMOS camera (Hamamatsu Photonics). Standard imaging media (extracellular fluid; ECF) consisting of 140 mM NaCl, 3 mM KCl, 1.5 mM CaCl2, 1 mM MgCl2, 5.5 mM glucose, 10 mM HEPES (pH 7.3), B27 (Gibco), glutamax (Gibco) was used, with 50 mM D-AP5 (Tocris; 0106), 20 mM CNQX (Tocris; 1045), and 100 mM picrotoxin (Tocris; 1128) added to block postsynaptic signals. Single image planes were acquired with 5.6 ms exposure (178.6 Hz) using Sedat 490/20 nm filter excitation and 525/36 nm filter emission. 400 frames (5.6 ms exposure/ 2240 ms total) were collected, starting 450 ms after the initial frame. A Model 4100 Isolated High Power Stimulator (A-M Systems; 930000) was used with platinum wires attached to a field stimulation chamber (Warner Instruments; RC-49MFSH), integrated by an Axon Digidata 1550B (Molecular Devices). Field stimulus voltage was set to the lowest that reliably produced presynaptic calcium transients in >95% presynaptic boutons using synaptophysin-GCaMP6f as a presynaptic calcium reporter. Experiments were performed at 33-34°C. Temperature and humidity were controlled by an Okolab incubation controller and chamber.

### Two-photon imaging screen in mouse cortex

AAV2/1 Viruses encoding iGluSnFR variants in the NGR display vector (a glycosylphosphatidylinositol (GPI) anchor) were generated by the Janelia Viral Core. The NGR vector was chosen based on prior data demonstrating excellent expression and membrane localization in culture^26,37^ and preliminary observations that it produced higher expression than the previously-used COBL9 GPI anchor. To better compare performance to a more conventional vector, we including iGluSnFR3 controls of both NGR and PDGFR vectors.

C57BL/6J male mice (Jackson Laboratory) of 2 months of age were anesthetized and placed in a stereotactic mount. Following craniotomy and dura removal, viral injections were done at two sites in the left visual cortex area V1 (−4.0 A/P, -2.8 M/L, 0.3 D/V and -2.5 A/P, same M/L and D/V coordinates), using a Nanoject III injector and a beveled borosilicate micropipette. Sparse expression necessary to distinguish individual neurites was achieved by injecting 250 nL of high-titer flexed indicator and low-titer Cre AAV mix (vg/mL): 2E12 AAV1-hSyn-Flex-<IGLUSNFR> + 1.2E8 AAV9-CamKII-0.4.Cre.SV40 (Addgene 105558-AAV9).

Two-photon microscopy imaging was done on a 12 kHz commercial resonant scanning microscope (Bergamo II, Thorlabs; ScanImage, MBF Bioscience) using an Olympus 25x 1.0 NA objective equipped with a spherical aberration correction collar to correct for the imaging window coverslip. Two-photon excitation was done at 1030 nm (Insight X3, Newport) using a fixed 46 mW power post-objective for all iGluSnFR variants and imaging depths. Fluorescence was collected using a GaAsP photomultiplier through a 525/50 band-pass filter (Semrock). For *in vivo* indicator screening, scans were acquired at 430 Hz, using 50 lines x 128 pixels/line at 10x software zoom, with scans lasting 2 minutes, while mice were headfixed, awake and visually stimulated with a 1 sec on/off diffuse light pulse that illuminated the microscope enclosure. Further functional characterization of select variants was done using moving gratings at different orientations in 45 deg increments.

Custom Matlab scripts were used to first motion correct raw scans using a cross-correlation template-maximization approach, followed by automated identification and extraction of iGluSnFR sources and events. In a first pass, the algorithm generated best-estimates for source localization based on fluorescence transients, which were clustered into larger patches. These initial source estimates were further refined using non-negative matrix factorization and a least-squares fit to yield source footprints from which fluorescence signals were extracted and further processed to characterize transients.

Several measures were used to compare indicator fluorescence properties: decay time constant, event signal-to-noise ratio (eSNR), event detectability, fluorescence skewness, bleaching factor and baseline fluorescence. On a source level, event decay time constant was calculated by first dividing the recorded fluorescence into 10 second windows, and within each window, the signal was cross-correlated at incremental time-lags up to a maximum lag followed by a robust linear model fit^48^ to the logarithm of the cross-correlated time-lags. To obtain the decay time-constant within each 10 second window, only statistically significant linear fits were considered (p<0.001), and was calculated as the absolute value of the inverse of the estimated slope. To reduce bias due to bursts that can obscure single-event fluorescence decay, a branch-level decay time-constant was calculated as the minimum time-constant across all identified sources from the same branch.

Spine turnover measurements were performed on an overlapping cohort of mice as the above screen, using the same surgical preparation. 3 mice received AAV1 encoding membrane-tagged GFP (hSyn-FLEX-EGFP.CAAX) in place of iGluSnFR, as control. Starting 2 weeks after injection, recordings were performed over 3 sessions (days 0,7,14) as for the activity screen. Mean motion-registered images from the recordings were analyzed to quantify spine turnover. Annotations were performed manually via a custom-written Python GUI, blinded to experimental condition.

### In vivo spatial specificity measurements

Two-photon activity imaging imaging was conducted with the same microscope and settings as in the previous section. To assess spatial specificity of the glutamate indicators, mice were presented with moving grating visual stimuli (2 Hz temporal frequency, 0.05 cycles/deg) of 8 directions (45 degree increments) in randomized order. Stimuli were presented for 2 sec following a 1 sec mean luminance stimulus. Each grating was presented 15 times during each recording session.

For each movie recorded in this way, ΔF=F-F0 was calculated for each pixel, where F0 is a moving median of F with a window of 10001 frames. Then, the “time-resolved tuning curve” was calculated as the average two second trace across trials of each orientation presentation. To get the “eight-point tuning curve,” the mean response over the 0.5 second before the stimulus was subtracted from the mean response during the on periods. Preferred orientation was calculated as the direction of the vector sum of response vectors, where each vector’s direction is the corresponding stimulus orientation and the vector magnitude is the response magnitude from the eight-point tuning curve. Orientation selectivity index (OSI) was calculated as the magnitude of the vector sum of response vectors. Response amplitude was calculated as the average of the magnitudes of response vectors. Pixelwise tuning maps were drawn using the preferred orientation as the hue, OSI as the saturation, and response amplitude as the value for each pixel.

To quantify degree of spatial specificity, cross-validated covariances were calculated for pairs of pixels. For each pair of pixels, the time-resolved tuning curve was calculated for each pixel using a mutually exclusive random split of trials, and the covariance between tuning curves was calculated. This was done over 20 random splits for each pair of pixels, and the resulting cross-validated covariance was the average of covariances over the 20 splits. This analysis was done separately on the set of labeled pixels (pixels on or near the brightest pixels) and background pixels (all other pixels).

The distance it took for the average cross-validated correlation across recordings to reach 1/e and 1/e2 of the maximum were recorded. Confidence intervals for these statistics were calculated by bootstrapping on recordings and using 2.5th and 97.5th percentile of the bootstrap distribution from 1000 bootstrap samples. The difference in statistic was deemed significantly different if the 95% bootstrap confidence interval of the difference in statistic did not include 0.

### In vivo AO2P imaging and data analysis

WT mice (C57BL/6J, male, 6-8 weeks old, Jackson Laboratory) were anesthetized with isoflurane using a precision vaporizer (3% v/v for induction and 1-2% v/v for maintenance) along with subcutaneous injection of buprenorphine (0.1 µg per body weight). Body temperature was maintained at 37 C with a heating pad during surgery. The animal was placed in a stereotaxic frame. A small hole (∼0.2-0.5 mm) was drilled at 1.7 mm posterior to the bregma and 1.8 mm lateral from the midline, corresponding to the transverse location of the barreloids of the right VPM nucleus. Cre-dependent iGluSnFR3.v857.GPI, iGluSnFR4s.PDGFR and iGluSnFR4f.PDGFR, were diluted and mixed to the final titer of 8×10e12 to 1.1×10e13 GC mL−1 for the GluSnFR virus and 3×10e10 for the Cre virus. The viruses were injected into the VPM at a depth of 3.25 mm and 3.1 mm at 1.6 mm posterior, 1.8 mm lateral and 1.65 mm posterior, 2.0 mm lateral at 50 nL, 10 nL min-1 at each location. A 4-mm craniotomy was performed over the right vS1 cortex (centroid at 1.6 mm posterior to the bregma and 3.3 mm lateral from the midline). A cranial window by a single 4-mm round coverslip (no. 1) was embedded in the craniotomy and sealed with cyanoacrylate glue (Loctite, catalog no. 401). Meta-bond (Parkell) was further applied around the edge to reinforce stability. A titanium head-bar was attached to the skull with Meta-bond and the remaining exposed bone was covered with dental acrylic (Lang Dental).

In vivo imaging was carried out after 4 weeks of expression and 3 days of habituation for head fixation. All imaging experiments used head-fixed awake mice under AO2P. AO correction was applied at the imaging depth below 350 µm^56^. For functional imaging, the laser was tuned to 960 nm, with a post-objective power of 80-100 mW. The frame rate was 240 Hz. The vibrissae of the mice were trimmed 3 days before functional experiments, leaving only two vibrissa among C1, C2, D1, D2, or Beta, whose corresponding cortical column had an optimal expression of iGluSnFR. The mice were head fixed to the imaging rig with a running disk. The whisker was illuminated with an infrared LED (M940L3, Thorlabs) and captured with a camera operated at 500 Hz and synchronized to the imaging setup.

Air-puff deflection was used for vibrissa stimulation. Pulse-controlled compressed air, 20-ms pulse width, 5 p.s.i. at the source, was delivered through a fine tube, which was placed parallel to the side of the mouse snout and 20 mm away from the targeted vibrissa. The frequency of the air puffs was from 2 to 30 Hz. The experiments were carried out in the absence of visible light. Within each barrel, eight regions (30×60 µm) were imaged at each stimulus frequency in random order to avoid any bias from photobleaching.

The frames in the raw two photon recordings a were motion corrected^57^ and background subtracted. The average and standard deviation of the frames were sent to a bouton mask extraction algorithm using adaptive thresholding and estimating the connected components that were within the expected size for individual boutons. The fluorescence signals of each bouton were linearly interpolateFree-whisking experiments were conducted by placing a thin pole on a stage capable of moving randomly in the rostro-caudal and medio-lateral directions, where the pole is occasionally within the reach of the mouse whiskers. The data were recorded while the mouse was in its exploratory phase, during which the whiskers swept rhythmically. The whisker location and structure along with the touch events, were identified using a maskless posture estimation package^58^.

### In vivo two-photon imaging and cell-attached recordings

Adult C57Bl/6 (8-16 weeks) mice were anesthetized with isoflurane (2% for induction and 1.5% for maintenance, in pure oxygen). The body temperature was continuously monitored and kept at 37.5 °C by a heating plate. Both eyes were covered with ophthalmic ointment. The skin above the left hemisphere was carefully removed by fine scissors to expose the skull following the administration of an analgesic (Metamizole, 200 mg/kg) and a local anesthetic (2% xylocaine). The exposed skull was left dry and cleaned with a sterile scalpel blade. A custom-made headplate was fixed to the cleaned skull with light-cured dental cement. For the chronic window, a 3 mm circular craniotomy was made above the left primary visual cortex (V1), and a matching coverslip was glued to the craniotomy using vetbond. For single-cell plasmid electroporation, the coverslip on the craniotomy had a small perforation that allowed the access of a patch-pipette to the cortical tissue. The procedure for single-cell electroporation was the same as previously described^26^. Briefly, plasmids encoding pAAV.hSyn.iGluSnFR variants in the PDGFR backbone were dissolved in an artificial intracellular solution to a concentration of ∼100 ng/ul and was delivered to random layer 2/3 neurons using a 4-5 MΩ patch-pipette. After electroporation, the perforated coverslip was carefully removed and replaced by an intact one. The hippocampal window was similarly prepared. After making a 3.0 mm craniotomy on the right hemisphere, the cortical tissue was carefully removed with a blunt needle that was connected to a vacuum pump. The corpus callosum and external capsule were also removed to expose the alveus of the hippocampus. The hippocampal window assembly (consisting of a coverslip and a stainless-steel cannula) was implanted so the coverslip was directly on top of the hippocampus.Two-photon glutamate imaging was performed with a custom-built two-photon microscope based on a commercial upright microscope (Slicescope, Scientifica). The microscope featured a 12-kHz resonant scanner (Cambridge Technology) and a tunable Ti:sapphire laser (Mai Tai HP DeepSee, Spectra-Physics). Microscope control and data acquisition were achieved using a custom LabVIEW GUI that supported a frame rate of up to 500 Hz. The laser was tuned to 950 nm for glutamate imaging and the power under the objective (40x, NA 0.8, Nikon) was kept below 30 mW to prevent photodamage.

Dendrites of L2/3 pyramidal neurons in mouse V1 were imaged at 200 Hz frame rate and glutamate signals from individual synapses were probed using full-field square wave drifting gratings (0.03 cycle per degree spatial frequency, 2 Hz temporal frequency, 2 s duration, 8 directions). Drifting gratings stimuli were generated using a custom Android app and presented on a tablet (Samsung, 10.5-inch) equipped with an OLED display. The tablet was placed 10 cm in front of the right eye of the mouse, covering a visual space of 96 degrees in azimuth and 70 degrees in elevation, and had a mean brightness of 4.8 cd/cm2.

For simultaneous two-photon axonal imaging and cell-attached recordings, the coverslip was changed to one with an access opening and warm artificial cerebro-spinal fluid (ACSF) (125 mM NaCl, 26 mM NaHCO_3_, 4.5 mM KCl, 2 mM CaCl_2_, 1.25 mM NaH_2_PO_4_, 1 mM MgCl_2_ and 20 mM glucose) was perfused throughout the experiment to keep the brain temperature constant. Loose seal cell-attached recordings from glutamate sensors expressing cells were performed with a patch-pipette filled with ACSF containing 25 μM Alexa Fluor 594. Axonal boutons from the same neuron were identified and imaged at 500 Hz framerate. Electrophysiological signals were acquired in either current-clamp mode or voltage-clamp mode by a patch-clamp amplifier (EPC10, HEKA).

All data were processed using custom scripts in MATLAB, and statistical tests were conducted in R. ROIs were automatically segmented or manually drawn based on the local correlation image. The Mean fluorescence signal F from each ROI was extracted, and the fluorescence change was calculated as ΔF/F = (F-F_0_)/F_0_. F_0_ was the mean of baseline fluorescence. All traces were denoised using a first-order Savitzky-Golay filter (25 ms window) except for the kinetics analysis.

To quantify the kinetics of each variant, spike-triggered averages of single-AP-associated glutamate signals were calculated for each axonal bouton. Because of the much slower time course of iGluSnFR4s, only single APs separated by more than 500 ms from the preceding and following APs were selected. The resulting single-AP glutamate transients were fitted to the following exponential equation.

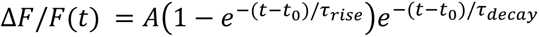

Where A is the amplitude factor, t_0_ is the time of the AP, and τ_rise_ and τ_decay_ are time constants for rise and decay. The amplitude was measured as the peak of the fitted trace, and time-to-peak was the time between the spike and the peak.

To compare the two iGluSnFR4 variants in resolving synaptic orientation selectivity in the visual cortex, the orientation tuning curve for individual synapses was obtained by calculating either the amplitude or the mean of the response. The tuning curve was then mapped onto orientation vector space and summed. The preferred orientation was the angle of the resultant vector. Orientation selectivity index (OSI) was calculated as the length of the resultant vector divided by the sum of responses to each orientation. The tuning magnitude was defined as the 2-norm of the tuning curve. For Pixelwise orientation maps, the hue is the preferred orientation (normalized to 180 °), the saturation corresponds to OSI, and the lightness is the tuning magnitude (normalized to the maximum of each field of view). To compare recordings from the hippocampus with data from V1, the denoised ΔF/F trace was converted to a modified z-score trace using the mean and the standard deviation from a baseline period. The baseline period is a 0.5 s period with the lowest standard deviation. Spontaneous glutamate transients were automatically detected using a threshold of 8 times the baseline standard deviation.

### Retinotopic mapping of L5 neuron dendrites

Experiments were performed on six wild-type mice (C57Bl6/J), aged between 4–6 weeks, of both sexes. Animals were anaesthetised with isoflurane (1-2% in Oxygen), their body temperature was monitored and kept at 37-38°C using a closed-loop heating pad, and the eyes were protected with ophthalmic gel (Viscotears Liquid Gel, Alcon Inc.). Analgesic (Rimadyl, 5 mg/kg) was administered subcutaneously before the procedure, and orally on subsequent days. Dexamethasone (0.5 mg/kg, IM) was administered intramuscularly 30 min prior to the procedure to prevent brain edema. The surgery began with the implant of a head-plate over the right hemisphere of the cranium. The head was shaved and disinfected; the cranium was exposed and covered with biocompatible cyanoacrylate glue (Vetbond, 3M). A stainless-steel head plate with a 10 mm circular opening was secured over the skull using dental cement (Super-Bond C&B, 10 Sun Medical Co. Ltd., Japan). Then a 3 mm wide square craniotomy was opened over V1 (centred at -3.3 mm AP, 2.8 ML from bregma). The exposed brain was constantly perfused with artificial cerebrospinal fluid (150 mM NaCl, 2.5 mM KCl, 10 mM HEPES, 2 mM CaCl2, 1 mM MgCl2; pH 7.3 adjusted with NaOH, 300 mOsm). We delivered 60 nL injections at multiple sites, at a cortical depth of ∼500 µm, to virally express iGluSnFR variants in layer 5 pyramidal neurons. We achieved sparse expression by injecting diluted AAV1-CaMK2a-Cre (2 × 10⁸ GC /ml), mixed with a concentrated: AAV1 hSyn.FLEX.iGluSnFR3.v857.GPI (final concentration: 3.88 × 10¹² GC /ml; 3 mice); or pGP-AAV-hSyn.FLEX.iGluSnFR4.v8880.PDGFR (final concentration: 2.3 × 10¹² GC /ml; 2 mice). For other purposes, the injection mix included also AAV1.hSyn.FLEX.NES-jRGECO1a.WPRE.SV40 (final concentration: 2.2 × 10¹² GC /ml). Following the injections, the craniotomy was sealed with a glass cranial window, assembled from a circular cover glass (4 mm diameter, 100 µm thickness) glued to a smaller insert (3 mm wide, 300 µm thickness) with index-matched UV curing adhesive (Norland #61).

Recordings of neuronal activity were performed 1 month after surgery with a standard resonant-scanning two-photon microscope (Bergamo II, Thorlabs), equipped with a Nikon 16x, 0.8 NA objective. The microscope was controlled using ScanImage 2023.1 (Ref 1). Excitation light was provided by a femtosecond laser (Chameleon Discovery TPC, Coherent) at 950 nm. Laser power was depth-adjusted between 25-50 mW. Sample fluorescence was collected in the green (525/50 nm) band. During recordings, mice were head fixed over a metal mesh wheel, where they could run at will. For each FOV, a structural z-stack served as a reference to identify L5 neurons and trace their apical dendrites back to the surface. For each neuron, as many apical dendrites as possible were recorded in sequential and longitudinal acquisitions, at a cortical depth ranging from 10 µm to 100 µm below the brain surface. Functional imaging was performed at, scanning field of views of 512×512 pixels at 30Hz, with a resolution ranging from 0.1894 to 0.1088 µm per pixel.

Visual stimuli were generated in Matlab (MathWorks) and displayed on 3 gamma-corrected LCD monitors (Adafruit 1.8” TFT LCD Display, resolution 128x160 px) surrounding the mouse at 90 degrees to each other. The LCD screens were covered with Fresnel lenses to correct for viewing angle inhomogeneity of the LCD luminance. The mouse was positioned at the centre of the U-shaped monitor arrangement at 20 cm from all three monitors, so that the monitors spanned ±135 degrees of horizontal and ±35 degrees of the vertical visual field. Sparse, spatial white noise stimuli were used to map the retinotopy of the imaged area and estimate the receptive field (RF) of spine inputs. Patterns of sparse black and white squares (4.5-6 degrees of the visual field) on a grey background were presented at 5 Hz, typically in 10 min sequences repeated 3 times for each FOV. At any point in time, each square had a 2% probability of being non-grey, independent of the other squares.

### Analysis of retinotopic mapping data

Recordings were pre-processed using Suite2p^59^. The final selection of ROIs was curated manually to eliminate noisy ROIs and retain only dendritic branches from the target neuron in a given imaging session.

Linear neuronal spatiotemporal RFs were estimated by regularised linear regression between the neuronal responses and the stimulus history. The Laplacian of the receptive field (RF) in space was regularized to enforce smoothness. stRFs were modeled using time-lagged regression: the stimulus matrix was constructed as the Toeplitz matrix containing the stimulus history shifted at different lags between t=0 and t=1.4s. ON and OFF subfields were fit simultaneously using separate stimulus predictors for increases and decreases in luminance at the same pixel, respectively. The ON/OFF subfield is computed as the average of the ON and OFF subfields while considering their respective signs to maintain the correct polarity.

For each ROI, we used 3-fold cross-validation to choose the regularization parameter that maximized the stRF predictive performance. The RF performance was measured as the explained variance (EV), defined as:

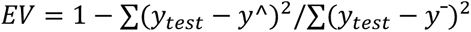

where *y*_test_ is the actual neuronal response, *y*^ is the predicted response, and *y*^-^ is the mean response. The final stRF was then fit with the best λ fixed, over the full recording.

We estimated stRF significance using a circular shift test. For each ROI, we fit the RF after circularly shifting the stimulus predictor in time, and repeated this procedure 1,000 times to build a null distribution of explained variances. A p-value was then calculated as the fraction of the explained variance from the null distribution that exceeded the explained variance from the real, unshifted data. ROIs were considered responsive if they met two criteria, 1) explained variance from the original dataset exceeded 0.1 and 2) the p-value from the circular shift test was smaller than 0.005.

The spatial RFs of responsive ROIs were estimated as the slice of the stRF bearing the largest response and were subsequently fit with a 2D Gaussian. This Gaussian fit represents a localized elliptical region where the ROI exhibits a response.

### Fiber photometry

Surgeries for in vivo fiber photometry recordings (Fig. 5) were performed as previously described^60^. In brief, we performed bilateral (100-150 nL each hemisphere, 2.0 × 10¹² GC /ml) injections of AAV (php.eB) encoding hSyn.FLEX.iGluSnFR variants into the VTA of Gad2-ires-Cre mice (JAX 028867). In each mouse, one random hemisphere received iGluSnFR4.PDGFR and the other either iGluSnFR3.v857.PDGFR or SF-iGluSnFR.A184S.PDGFR. The coordinates targeting VTA were (AP:-3.05; ML:±1.75; DV:-4.2 from pia; ±15 degree). After a minimum two-week recovery period following surgery, the mice underwent mild water restriction. We used a custom made CMOS-based fiber photometry system as previously described^61^. In brief, both GFP-based SF-iGluSnFR and YFP-based SF-iGluSnFR were excited with a 470 nm LED and emission signals were collected with a conventional GFP emission filter (520 ± 35 nm). CMOS sensors and LEDs were controlled by a Teensy 4.1 microcontroller and images from CMOS sensors were acquired using a custom program written in Bonsai^62^.

The same Bonsai instance was used to generate sequences of paired stimuli consisting of a tone (5 kHz pure-tone, ∼70 dB, 1 s duration) followed by a 1 s delay and a subsequent water reward (2 μL). Inter-trial intervals were randomized. To specifically examine reward responses independent of learned associations, we analyzed only the initial sessions before mice had developed associations between the tone and reward delivery.

Data were preprocessed and analyzed using custom Python scripts. Mice with significant motion artifacts were excluded from analysis. The time course of raw CMOS pixel values was detrended using a mild Gaussian low-cut filter (cutoff: 2–4 min) and corrected for slow baseline drift caused by photobleaching using a 4th-order polynomial fit. For generating peri-event time histograms (PETH; Fig. 5c), traces from individual trials were locally baseline-subtracted using the 2 s period preceding reward consumption, and then averaged across trials.

## Acknowledgements

We thank the Lab Animal Services and Neurosurgery & Behavior teams at the Allen Institute for technical support, and to Brooke Wynalda and Sujatha Narayan for assistance in cloning. We thank Tim Brown, the Tool Translation Team, and Deepika Walpita (Janelia Research Campus) for providing resources and access to instrumentation. We thank Ilme Schlichting for X-ray data collection, the European Synchrotron Radiation Facility (ESRF) for provision of synchrotron radiation facilities, and Estelle Mossou for assistance and support in using beamline ID23-1.

Work at the Allen Institute was funded by grants from the National Institutes of Health (NIH), specifically, 1DP2NS136990 and BRAIN Initiative UM1MH136462 to KP and 1F30MH138009 to MEX, by the Paul and Daisy Soros Fellowships for New Americans to MEX, by the Human Frontier Science Program HFSP(LT0052/2022-L) to KMH, and by the Allen Institute for Neural Dynamics. We thank the Allen Institute founder, P. G. Allen, for his vision, encouragement and support.

Work at Janelia research campus was funded by the Howard Hughes Medical Institute.

Work at TUM was funded by the Deutsche Forschungsgemeinschaft (KO 979/7-1) and the Max-Planck-School of Cognition. AK is a Hertie-Senior-Professor for Neuroscience.

Work at UCL and IIT was funded by the Armenise-Harvard Foundation (CDA to LFR), the Human Technopole (HT–ECF 3588 to LFR), the Boehringer Ingelheim Foundation (studentship to AL), the Biotechnology and Biological Sciences Research Council (studentship to AL), and by UKRI (Frontier Award EP/X022366/1 to MC).

Work at St. Jude Children’s Research Hospital was funded by the American Lebanese Syrian Associated Charities (ALSAC).

Work at UCSD was funded by grants from the National Institutes of Health (NIH), specifically, U19 NS137920, U24 EB028942, R01 NS143141, and U01 NS126054 to DK. RI is a Schmidt Science Fellow.

## Author contributions

**Conceptualization:** Abhi Aggarwal, Jeremy Hasseman, and Kaspar Podgorski.

**Data curation:** Abhi Aggarwal, Adrian Negrean, Yang Chen, Rishyashring Iyer, Daniel Reep, Anyi Liu, Michael E. Xie, Kenta M. Hagihara, Lucas W. Kinsey, Julianna L. Sun, Pantong Yao, Benjamin J. Arthur, and Kaspar Podgorski.

**Formal analysis:** Abhi Aggarwal, Adrian Negrean, Yang Chen, Rishyashring Iyer, Daniel Reep, Anyi Liu, Anirudh Palutla, Michael E. Xie, Kenta M. Hagihara, Lucas W. Kinsey, Julianna L. Sun, Jihong Zheng, Arthur Tsang, Benjamin J. Arthur, Miroslaw Tarnawski, Srinivas C. Turaga, Alison G. Tebo, L. F. Rossi, David Kleinfeld, and Kaspar Podgorski.

**Funding acquisition:** Anyi Liu, Matteo Carandini, L. F. Rossi, David Kleinfeld, Arthur Konnerth, Karel Svoboda, and Kaspar Podgorski.

**Investigation:** Abhi Aggarwal, Adrian Negrean, Yang Chen, Rishyashring Iyer, Daniel Reep, Anyi Liu, Michael E. Xie, Bryan MacLennan, Kenta M. Hagihara, Lucas W. Kinsey, Julianna L. Sun, Pantong Yao, Jihong Zheng, Arthur Tsang, Getahun Tsegaye, Ronak H. Patel, Julien Hiblot, Philipp Leippe, Miroslaw Tarnawski, Jonathan S. Marvin, L. F. Rossi, Jeremy Hasseman, and Kaspar Podgorski.

**Methodology:** Abhi Aggarwal, Adrian Negrean, Yang Chen, Rishyashring Iyer, Daniel Reep, Michael E. Xie, Kenta M. Hagihara, Jihong Zheng, Arthur Tsang, Getahun Tsegaye, Julien Hiblot, Philipp Leippe, Miroslaw Tarnawski, Srinivas C. Turaga, Alison G. Tebo, Matteo Carandini, L. F. Rossi, David Kleinfeld, and Kaspar Podgorski.

**Project administration:** Abhi Aggarwal, Julien Hiblot, Srinivas C. Turaga, Alison G. Tebo, David Kleinfeld, Arthur Konnerth, Glenn C. Turner, Jeremy Hasseman, and Kaspar Podgorski.

**Resources:** Daniel Reep, Julien Hiblot, Miroslaw Tarnawski, Jason D. Vevea, Srinivas C. Turaga, Alison G. Tebo, Matteo Carandini, David Kleinfeld, Karel Svoboda, Arthur Konnerth, Jeremy Hasseman, and Kaspar Podgorski.

**Software:** Yang Chen, Daniel Reep, Michael E. Xie, Kenta M. Hagihara, Lucas W. Kinsey, Benjamin J. Arthur, and Kaspar Podgorski.

**Supervision:** Abhi Aggarwal, Julien Hiblot, Jason D. Vevea, Alison G. Tebo, Matteo Carandini, L. F. Rossi, David Kleinfeld, Arthur Konnerth, Karel Svoboda, Glenn C. Turner, Jeremy Hasseman, and Kaspar Podgorski.

**Validation:** Abhi Aggarwal, Adrian Negrean, Yang Chen, Rishyashring Iyer, Daniel Reep, Michael E. Xie, Kenta M. Hagihara, and Kaspar Podgorski.

**Visualization:** Abhi Aggarwal, Adrian Negrean, Yang Chen, Rishyashring Iyer, Daniel Reep, Anirudh Palutla, Michael E. Xie, Kenta M. Hagihara, Lucas W. Kinsey, Julianna L. Sun, L. F. Rossi, and Kaspar Podgorski.

**Writing - original draft:** Abhi Aggarwal, Adrian Negrean, Yang Chen, Rishyashring Iyer, Anyi Liu, Michael E. Xie, Kenta M. Hagihara, Miroslaw Tarnawski, Srinivas C. Turaga, Alison G. Tebo, L. F. Rossi, David Kleinfeld, Arthur Konnerth, Glenn C. Turner, and Kaspar Podgorski.

**Writing - review & editing:** Abhi Aggarwal, Karel Svoboda, Glenn C. Turner, Jeremy Hasseman, and Kaspar Podgorski.

## Notes

### Competing Interest Statement

The authors have declared no competing interest.

### Summary of Updates

Methods updated; higher resolution figures used; stylistic changes (fonts and font sizes)

